# Strategies to decipher neuron identity from extracellular recordings in the cerebellum of behaving non-human primates

**DOI:** 10.1101/2025.01.29.634860

**Authors:** David J. Herzfeld, Nathan J. Hall, Stephen G. Lisberger

**Affiliations:** Department of Neurobiology, Duke University School of Medicine, Durham, NC, 27710, USA

**Keywords:** cell type, classification, Golgi cell, Purkinje cell, unipolar brush cell, mossy fiber, molecular layer interneuron

## Abstract

Identification of neuron type is critical to understand computation in neural circuits through extracellular recordings in awake, behaving animal subjects. Yet, modern recording probes have limited power to resolve neuron type. Here, we leverage the well-characterized architecture of the cerebellar circuit to perform expert identification of neuron type from extracellular recordings in behaving non-human primates. Using deep-learning classifiers we evaluate the information contained in readily accessible extracellular features for neuron identification. Waveform, discharge statistics, anatomical layer, and functional interactions each can inform neuron labels for a sizable fraction of cerebellar units. Together, as inputs to a deep-learning classifier, the features perform even better. Our tools and methodologies, validated during smooth pursuit eye movements in the cerebellar floccular complex of awake behaving monkeys, can guide expert identification of neuron type during cerebellar-dependent tasks in behaving animals across species. They lay the groundwork for characterization of information processing in the cerebellar cortex.

**Impact statement:** To understand how the brain performs computations in the service of behavior, we develop methods to link neuron type to functional activity within well-characterized neural circuits. Here, we show how features derived from extracellular recordings provide complementary information to disambiguate neuron identity in the cerebellar cortex.

## Introduction

Our goal is to understand how neural circuits generate behavior in awake, behaving monkeys by recording the extracellular activity of participating neural populations during carefully contrived behaviors^1,2^. Within any given neural circuit, different neurons feature different molecular, anatomical, connectional, and functional properties^3–11^. Thus, analysis of the coordinated processing by multiple, distinct neuron classes will be necessary to reveal the computational organization of the brain^11,12^. Yet, our main tool for studying how the neural circuits generate behavior, extracellular recording, is poorly suited to identification of neuron type.

We and many other laboratories now use multi-contact probes in both monkeys and mice, with the shared ability to recording from more than a few units simultaneously^13^ but the shared weakness that extracellular recordings offer poor access to identification of neuron type.

Optogenetic identification of neuron type is not an ideal solution. It is feasible (but fraught with challenges^14^) in awake mice, but normally gives access to a single cell type in a given preparation. Even under ideal conditions, genetic tagging of multiple specific cell types is limited by the overlapping spectral activation functions^15,16^ of a limited number of opsins and the limited genetic accessibility beyond rodents^17^. Here, we develop a strategy that allows us to cluster and label different neuron types recorded in a single circuit during sensorimotor behavior.

In the structure we study, the cerebellum, the classical circuit diagram (Figure 1A) includes multiple neuron types^18^ and two distinct groups of input fibers. Owing to their distinct extracellular signatures, neurophysiologists have focused primarily on the mossy fibers^19–24^ and climbing fibers^25,26^ that provide the main inputs to the cerebellum as well as one neuron type: the Purkinje cells^13,21,25–30^ that form the only output from the cerebellar cortex. Other neurons that likely perform critical computations inside the circuit have been comparatively ignored, including granule cells, unipolar brush cells, Golgi cells, and molecular layer interneurons. Thus, standard approaches address the question of how the output from the cerebellar cortex contributes to behavior in a few model systems, but not how the circuit works, what it computes, or how it transforms mossy fiber and climbing fiber inputs into Purkinje cell outputs. The strategies we explore here largely overcome that limitation by allowing direct mapping from features of extracellular recordings onto the identify of a neural unit.

**Figure 1.**
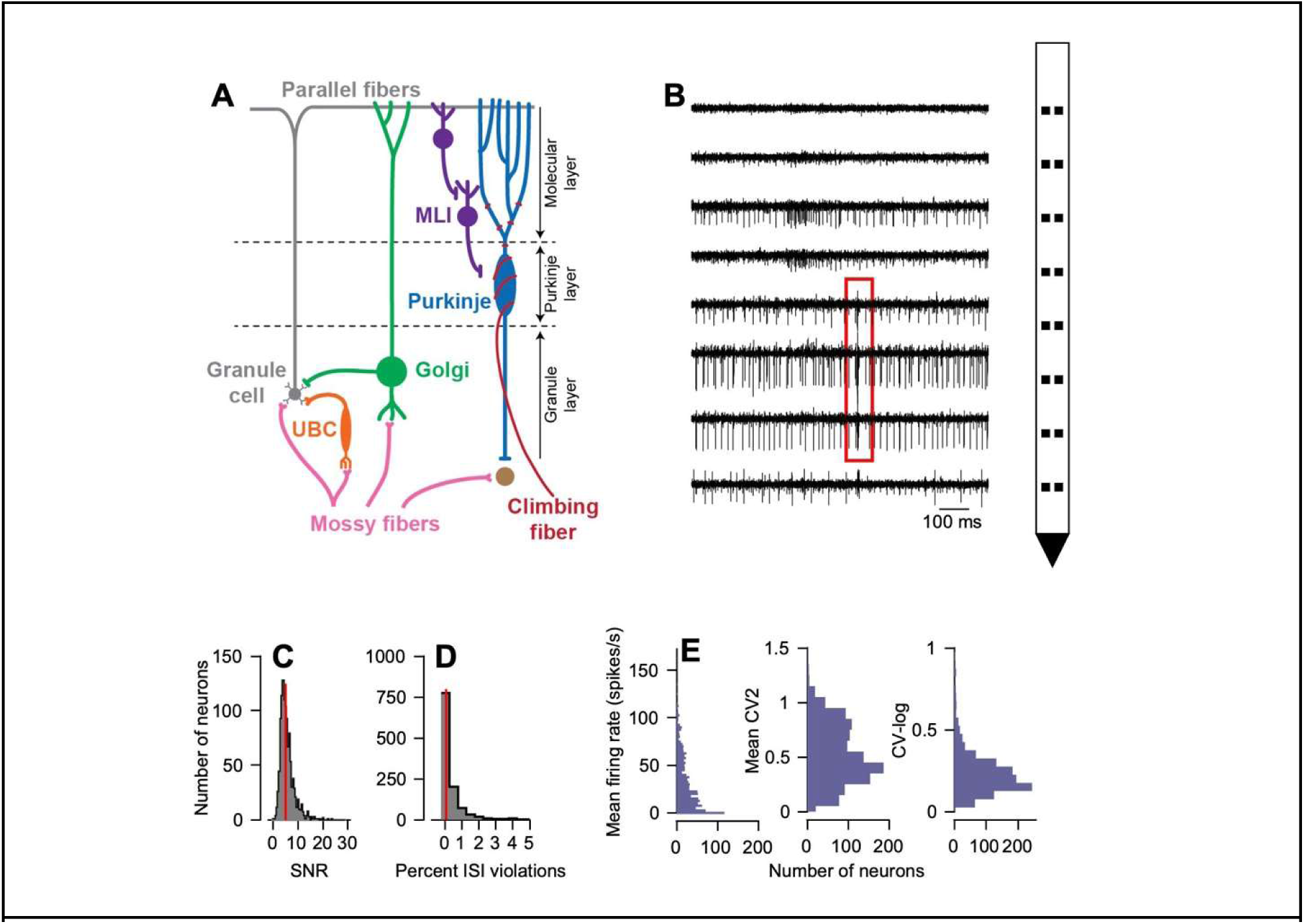
Properties of the neurophysiological recordings used to identify cerebellar neurons from monkeys. (A) Diagram of the canonical cerebellar circuit, simplified. (B) Exemplar recording from the floccular complex using a 16-contact Plexon S-Probe; the 8 traces show electrical activity on 8 channels in one column. Red box denotes an identified complex spike across contacts. (C) Distribution of signal-to-noise ratios across our full sample of neurons computed on the primary channel. (D) Distribution of the percent of spikes that occur within an assumed absolute refractory period of 1 ms across our full sample. Red lines in C-D denote the mean across all recorded units. (E) Distributions of scalar firing rate statistics of *n=1,152* recorded units shown as histograms. Left: the mean firing rate computed across each complete recording session. Middle: the mean CV2^36^. Right: the log of the coefficient of variation^32,33^.

In a multi-lab collaboration^14^, we and others showed recently that deep-learning neural networks can use features derived from high-density extracellular recordings to disambiguate ground-truth cerebellar cell types identified via optogenetic stimulation^31^. Now, we extend the previous study to allow neuron-type identification from features alone, without optogenetic stimulation. *<u>First</u>*, we provide tools and methods for expert labeling of neuron types from extracellular recordings in behaving non-human primates where viral tools for ground-truth labeling are still in their infancy^17^. The strong correspondence between our expert labels and the predictions of the ground-truth classifier validates our labeling approach^14^. *<u>Second</u>*, we extend the ground-truth classifier by exploring whether and how well we can inform neuron identity by high-dimensional features that are measured readily from extracellular recordings, in contrast to previous cerebellar cell-type classifiers that relied on scalar metrics^32–34^.

Our strategy uses data recorded from the cerebellar floccular complex during smooth pursuit eye movements. Using deep-learning approaches, we test the informativeness of 4 features for neuron identification: classical auto-correlograms; “3D” auto-correlograms that normalize for behaviorally-driven fluctuations in firing rate; the complete time course of waveform; and the spike-triggered LFP as an index of the local electrical environment. Each electrophysiological feature separately provides impressive information about neuron type, but as expected, the best classification performance is achieved by a classifier that uses multiple features. We hope that the next steps would: deploy the identification of neuron type to reveal circuit operation; use multiple electrophysiological features to identify additional cerebellar neuron types; and possibly implement a similar strategy in other brain regions.

## Results

The cerebellar circuit is composed of discrete neuron types^3,18^ arranged in a relatively uniform cytoarchitecture (Figure 1A). We can think of the circuit as performing a computation that transforms the cerebellar input signals from mossy fibers into the output from Purkinje cells. As a field, we already know how to identify recordings from mossy fibers and Purkinje cells in non-human primates^19,20,35^. Now, our goal was to provide an objective, quantitative basis for establishing the neuron types of the other single units recorded extracellularly from the cerebellar circuit using high-density probes in awake, behaving, non-human primates. We want to enable analysis of how circuits compute by providing a validated platform for neuron-type identification that goes beyond previous efforts.

Our challenge was to identify neuron type by taking advantage of the information available from knowledge of the cerebellar connectome (Figure 1A) and gleaned from extracellular recordings (Figure 1B). Our strategy was to record from the floccular complex of rhesus macaques during a behavior controlled by this part of the cerebellum^37^, smooth pursuit eye movements. Recent efforts^14,38–40^ suggest that we could discern neuron type from a combination of the statistics of discharge patterns, the shape of extracellular action potentials or their distribution across contacts, and the local electrical properties near the recordings. A critical first criterion for success is that spike-sorting, required for essentially all extracellular recordings with multi-contact probes, delivers well isolated single-units. We ensured excellent isolation by manual curation of the sorter’s output to ensure that all units had a high signal-to-noise ratio (mean ± SD, 5.6 ± 2.9, Figure 1C) and minimal violations of a 1-ms refractory period (mean number of violations, 0.6 ± 2.5%, Figure 1D).

The discrete metrics used in previous studies to automatically label cerebellar neuron types^32,33^ failed when applied to our recordings. Distributions of firing rate and discharge regularity did not show multiple peaks in their distributions that could have indicated potential heterogeneity of metrics across cell types. The distribution of mean firing rates across our population (Figure 1D, left) was broad (SD=28 spikes/sec) and unimodal (Hardigan’s dip test, D=0.008, p=0.95). “CV2” (Figure 1D, middle), a metric of discharge regularity that has been used previously to identify cerebellar cell types^33^, shows at most a hint of a non-significant (D=0.01, p=0.43) multi-modal distribution. We note that the majority of neurons had firing rate patterns that were more regular than Poisson (mean CV2=0.52 ± 0.28). Finally, the logarithm of the coefficient of variation (CV-log) across our sample (Figure 1D, right), a metric used previously to disambiguate cerebellar cell types^34^, revealed no evidence of a multimodal distribution (D=0.008, p=0.93).

### Ground-truth recordings from cerebellar Purkinje cells

A subset of Purkinje cells can be identified definitively, using either single electrodes or multi-contact probes, through simultaneous recording of their simple and complex spikes. In an example recording of a Purkinje cell (Figure 2A), we aligned individual voltage traces to the onset of *n=250* complex spikes (black arrow). We summarize the quintessential pause in simple spike activity following the occurrence of a complex^41,42^ in a complex-spike-triggered cross-correlogram (Figure 2B, top). In our sample of 111 ground-truth Purkinje cells, the duration of complex-spike-induced simple spike pauses ranged mostly from 10 to 25 ms, with a few longer pauses (Figure 2C).

**Figure 2.**
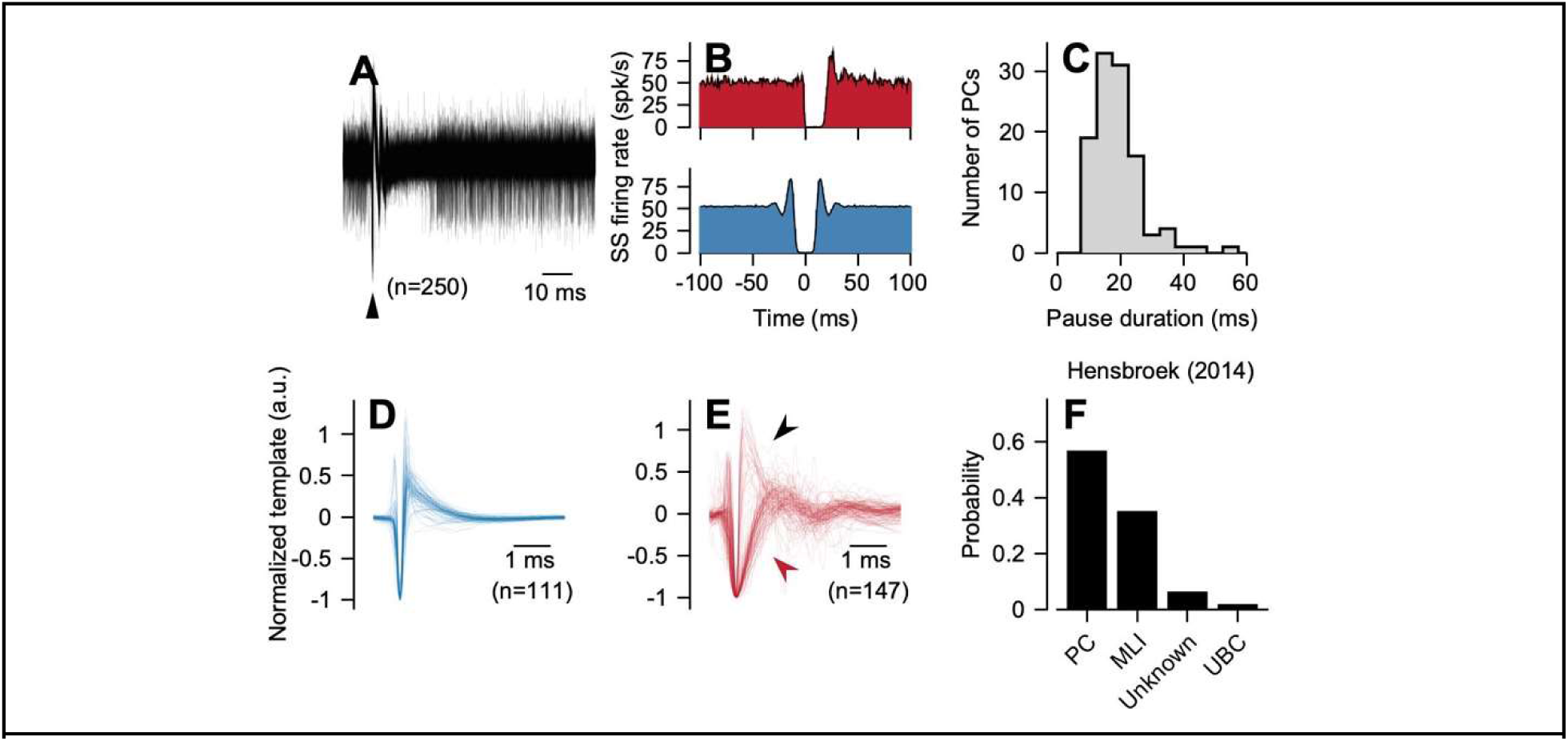
Firing rate properties of ground-truth identified Purkinje cell simple and complex spikes. (A) Example recording of a Purkinje cell’s simple spikes and complex spikes aligned to the 250 random occurrences of a complex spike (black arrowhead). Note the complex-spike-induced pause in the Purkinje cell’s simple spikes. (B) Simple spike cross-correlogram aligned to the occurrence of a complex spike at *t=0* ms (top, red) and simple spike auto-correlogram (blue, bottom), both for the Purkinje cell shown in (A). (C) Distribution of the duration of complex-spike-induced simple spike pauses across *n=111* ground-truth Purkinje cells. (D) Primary channel waveforms of ground-truth Purkinje cell simple spikes, normalized. (E) Same as (D) except for ground-truth Purkinje cell complex spikes. Black arrowhead points to presumed somatic complex spikes whereas red arrowhead points to dendritic complex spikes. (F) Probability of cell-type labels generated by a previously established classification algorithm^32^ when supplied with ground-truth Purkinje cell simple spikes from our data as input.

Other properties of ground-truth Purkinje cells were consistent across our sample. We will show later in summary graphs that the statistics of firing rate, as assessed by construction of auto-correlograms (ACGs, Figure 2B, bottom), were similar across ground-truth Purkinje cells. We also observed consistency in the simple spike action potential waveform as measured on the contact with the largest potential (Figure 2D). To allow comparison of the waveform shape across neurons, we normalized each waveform to its peak and reflected it, if necessary, so that the first major deflection always was negative. The primary channel waveform of complex spikes (Figure 2E) divided into two classes, with impressive uniformity within classes. Broad waveforms likely correspond to calcium spikes in the distal dendrites^43^ (red arrow); waveforms that show discrete spikelets (black arrow) likely correspond to post-synaptic climbing fiber responses recorded at or near the Purkinje cell soma^43^.

Given the ability to identify and characterize ground-truth Purkinje cells, we were able to test how well a previous cerebellar cell type classification algorithm^32^ would generalize to data from awake, behaving, non-human primates. The prior study showed excellent classification of recordings from Purkinje cells in awake rabbits (86% accuracy) based on mean firing rate, local firing rate regularity assessed via CV2, and the median absolute deviation of interspike intervals from the median. For our sample of ground-truth Purkinje cells, the previous algorithm classified only 57% (63/111) correctly as Purkinje cells (Figure 2F). The majority of incorrectly classified Purkinje cells were assigned by the other criteria in the classifier as molecular layer interneurons (39/111).

### Strategy

We develop our strategy for expert-identification of neuron type in five steps. 1) We leverage identification of the Purkinje cell layer to devise a quantitative approach to assign layers to different contacts on the probes. 2) We use layer information and cross-correlograms to identify molecular layer interneurons. 3) We develop quantitative criteria to divide neurons recorded in the granule cell layer into Golgi cells, unipolar brush cells, and mossy fibers. 4) We use deep-learning to test the informativeness of multiple, individual features of cerebellar recordings for neuron-type identification. 5) We demonstrate that a classifier based on the combination of multiple features yields neuron-type identification that agrees well with the ground-truth and expert assessments available to us in primates.

### Layer identification anchored by ground-truth Purkinje cells

The cerebellar cortex is a laminar structure: different neuron types with different electrophysiological signatures reside in different layers. The somas of Purkinje cells form a monolayer, termed the Purkinje cell layer, which can be identified in many multi-contact recordings by the presence of ground-truth Purkinje cells and transitions of complex spike waveforms from dendritic complex spikes recorded in the molecular layer to discrete spikelets recorded nearer the Purkinje cell soma^43^. Because recordings do not always yield a ground-truth Purkinje cell, we looked for discrete electrical signatures that could demarcate layers in the absence of a ground-truth Purkinje cell. For instance, previous work in the cerebral cortex has used current source density analysis derived from local field potentials (LFPs) to identify cortical layers^44–50^. In addition, a prior report showed consistent current source density profiles across layers from records in the cerebellum of anesthetized rats^51^, suggesting that layer-dependent signatures in the current source density analysis might extend to behaving cerebellar preparations.

To demonstrate that LFPs could establish cerebellar layer, we aligned our current source density analysis to the onset of smooth target motion in discrete trials, a sensorimotor stimulus known to strongly drive activity in the floccular complex^13,27,29,30^. The magnitude of the current source density shows a clear pattern across the depth of the electrode in an exemplar recording (Figure 3A). Direct comparison with the depth of the maximum-amplitude complex and simple spike waveforms of a ground-truth Purkinje cell recorded in the same session (Figure 3B, arrows; Figure 3A horizontal dashed lines) links the current source density profile to cerebellar layers. The same pattern of sources and sinks appears in the mean current source density map computed across all recordings with a ground-truth Purkinje cell (Figure 3C). Here, we aligned each recording to the electrode contact with the largest simple spike amplitude, corresponding to our estimate of the Purkinje cell soma. We reflected the current source density map computed for each recording, if necessary, to ensure that the primary complex spike channel was located at the top of the map. The stereotypical current source density profile in Figure 3C allows layer identification even in the absence of a ground-truth Purkinje cell in the associated recording.

**Figure 3.**
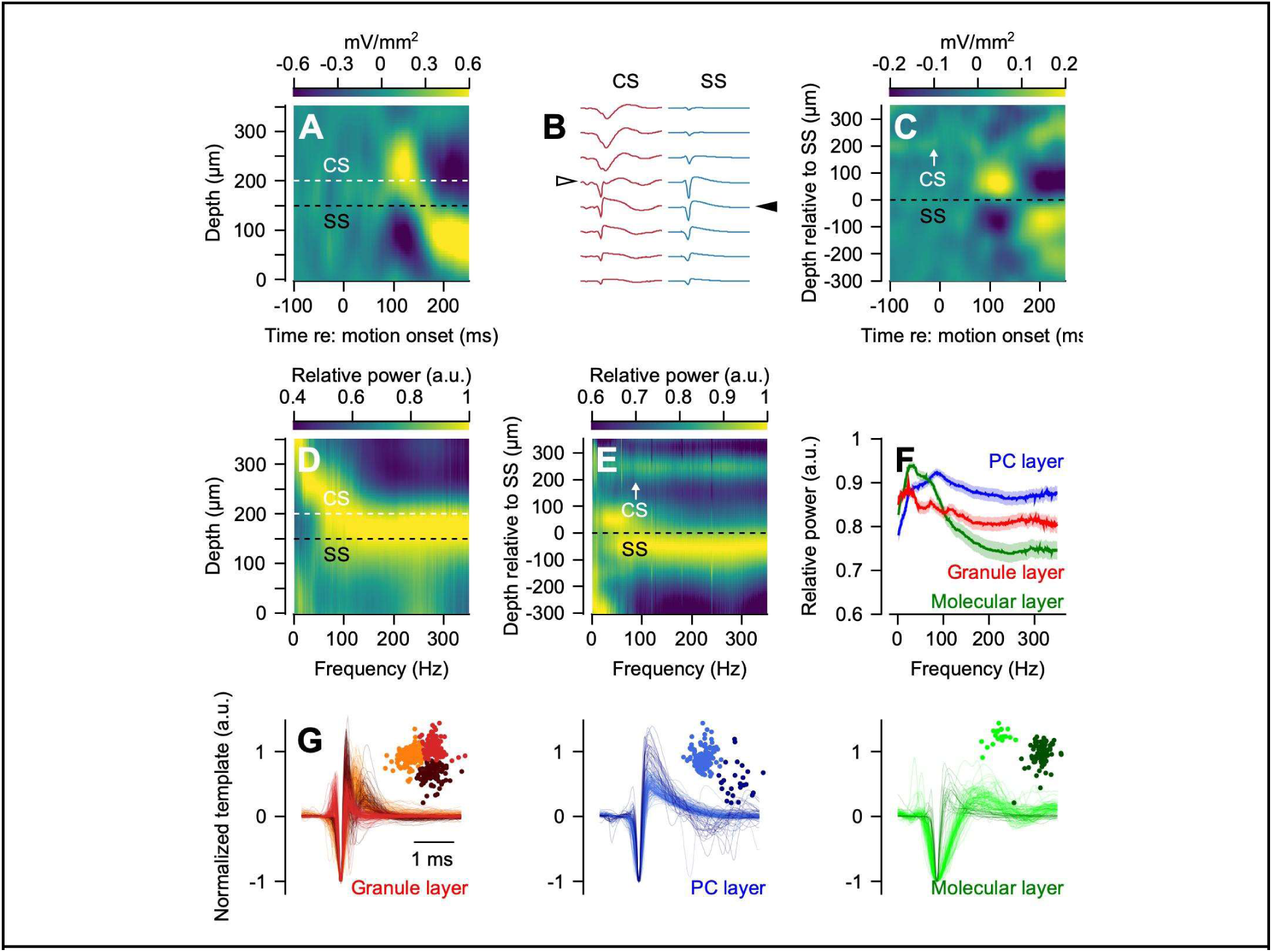
Identification of cerebellar layer from extracellular recordings. (A) Current source density computed from the local field potential for an example recording session. Horizontal lines denote the depth relative to the tip of the probe for the primary contact of simple spikes (black) and complex spikes (white) for a ground-truth Purkinje cell. (B) Complex spikes (red, left) and simple spikes (blue, right) for the ground-truth Purkinje cell recorded in (A) across contacts. Vertical position of each trace corresponds to the depth axis in (A). Arrowheads show the primary channel for the complex (white) and simple spikes (black). (C) Mean current source density across all recordings with a ground-truth Purkinje cell. Current source density maps were aligned with the primary channel of Purkinje cell simple spikes at a relative depth of 0 μm across recordings. Each recording was reflected about the 0 μm axis, if necessary, to ensure that the primary channel of the Purkinje cell complex spike had a positive value of relative depth. (D) Relative power of the LFP as a function of frequency across channels for the same recording from A-B. (E) Relative power of the LFP across channels, averaged across recording sessions with a ground-truth Purkinje cell. Preprocessing was performed as in (C). (F) Relative power of the LFP computed on primary contacts identified in the granule, Purkinje cell, and molecular layers. (G) Primary channel waveforms recorded in the identified granule (left), Purkinje cell (middle), and molecular (right) layers. We used K-means clustering following principal component analysis to split each layer’s waveforms into clusters (insets).

To ask whether the same approach could work in areas of the cerebellar cortex where the neuron-behavior relationship is unknown or the behavior isn’t structured into trials, we tested an alternative strategy to identify cerebellar layers using current source density analysis. A recent study in the cerebral cortex demonstrated that normalizing the LFP response across electrode contacts was sufficient to disambiguate layer^52^. We found complementary results in the cerebellar cortex. In a map of normalized frequency content across electrode contacts using the same recording as Figure 3A-B (Figure 3D), the Purkinje cell layer shows strong power in the upper frequency bands (50-350 Hz). The characteristic response persisted in averages across recordings with our full sample of ground-truth Purkinje cells (Figure 3E) when we aligned the depth based on the location of the contact with the largest simple spike waveform using the same convention as Figure 3C. Finally, averages of the normalized frequency spectra within layers identified manually revealed that the responses were substantially different for each layer (Figure 3G). We conclude that our use of local field potentials can generalize beyond situations such as in Figures 3A-C to establish the cerebellar layer of each recording contact. It is not necessary to align each current source density to the onset of the stimulus for eye movement or another behavior, or to collect data in trials.

Layer identification proved qualitatively useful for cell-type identification. We used unsupervised methods to cluster the action potential waveforms recorded in each layer identified through the analysis of the local field potentials (Figure 3G). Within each of the 3 layers, waveforms segregated into discrete clusters, shown by the different colored symbols in a space defined by the first two principal components of the waveforms (inset). Inspection of the waveforms reveals that they differed qualitatively between layers, as well as between clusters. The success of the qualitative analysis in Figure 3G encouraged us that information about waveform and layer would make major contributions to neuron-type identification.

### Functional identification of molecular layer interneurons

We identified molecular layer interneurons by their location in the molecular layer, a criterion used previously^53^, as established by normalized LFP and current source density profiles. For a subset of non-Purkinje cells recorded in the molecular layer, we were able to document an inhibitory connection at monosynaptic latencies to a simultaneously recorded ground-truth Purkinje cell. In the example pair illustrated in Figure 4A, we aligned simultaneously-recorded voltage traces of a putative molecular layer interneuron (top) and a nearby Purkinje cell (bottom) to 250 randomly selected spikes of the putative molecular layer interneuron (arrowhead). There is a noticeable reduction in the density of Purkinje cell simple spikes (Figure 4A, bottom) in the milliseconds following a spike in the putative molecular layer interneuron. We demonstrated inhibition from the molecular layer interneuron with a cross-correlogram of simple spike responses for the Purkinje cell aligned to the occurrence of spikes in the putative molecular layer interneuron (Figure 4B, bottom). We confirmed the quality of the isolation of the putative molecular layer interneuron with an auto-correlogram (Figure 4B, top) and the ground-truth identify of the Purkinje cell through a complex-spike triggered cross-correlogram of simple spike firing (Figure 4B, middle).

**Figure 4.**
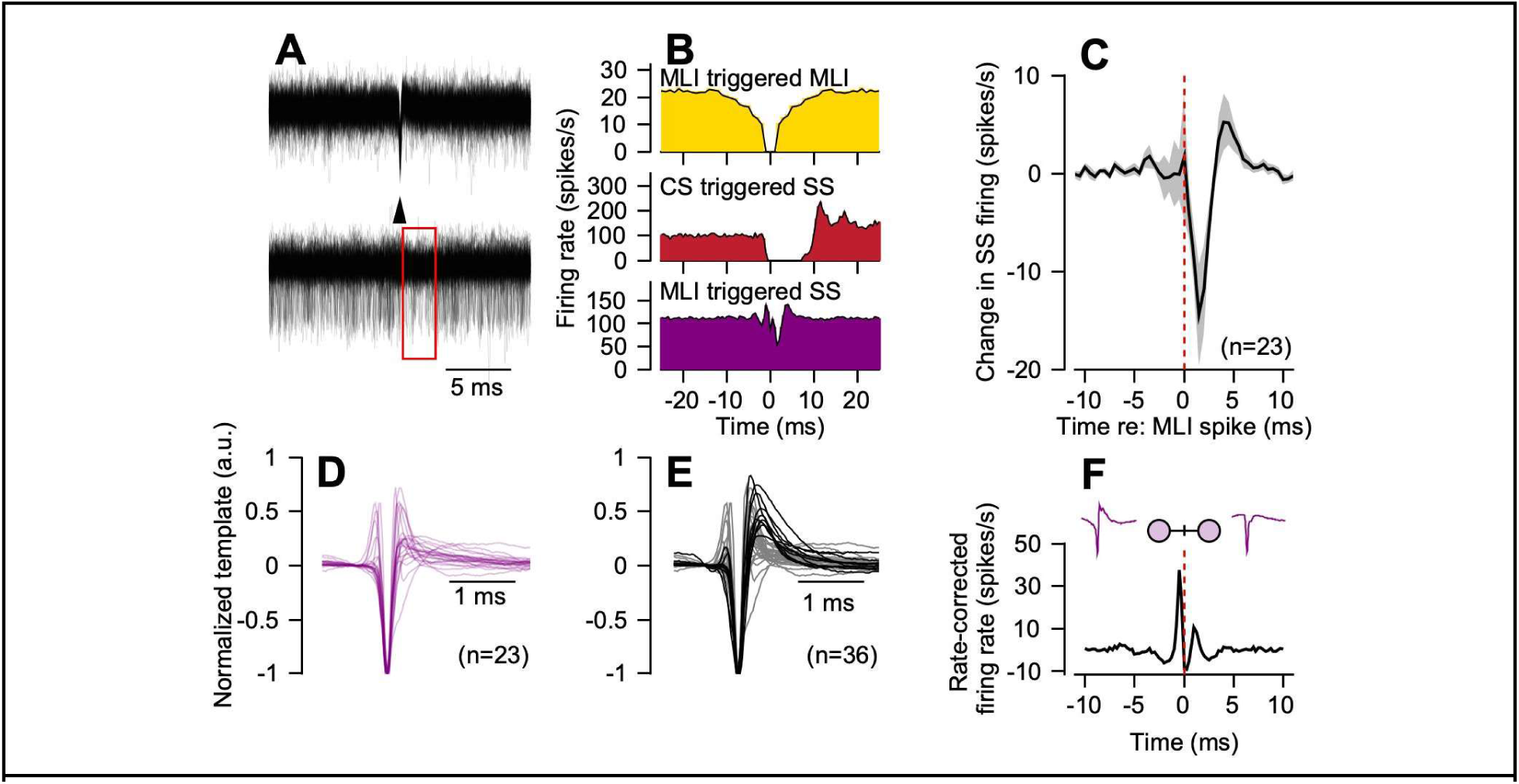
Functional identification of molecular layer interneurons. (A) Example recording aligned to the time of spikes in a functionally-identified putative molecular layer interneuron. Top: superimposed traces from the molecular layer interneuron’s primary channel, aligned at the arrowhead to *n*=250 randomly selected spikes. Bottom: the simultaneously-recorded primary channel of a ground-truth Purkinje cell’s simple spikes aligned to the same spikes and time points as the top plot. Note the subtle decrease in density of Purkinje cell simple spikes following the occurrence of a spike in the molecular layer neuron. (B) Top: auto-correlogram for the molecular layer interneuron in (A). Middle: simple spike cross-correlogram aligned to the time of a complex spike for the ground-truth Purkinje cell shown in (A). Bottom: simple spike cross-correlogram aligned to the time of a spike in the functionally identified molecular layer interneuron. (C) Mean cross-correlogram across 23 paired recordings showing the change from baseline of ground-truth Purkinje cell simple spike firing rates, aligned to the time of a simultaneously recorded putative molecular layer interneuron spike. (D) Normalized primary channel waveform for functionally identified molecular layer interneurons. (E) Primary channel waveforms of molecular layer interneurons identified functionally via their interaction with ground-truth Purkinje cells or their presence in the molecular layer. Waveforms shown in (D) are a subset of those in (E). Grey and black waveforms show the results of splitting the full sample based on hierarchical clustering into two groups with different typical waveform profiles. (F) Evidence for gap-junction coupling between a pair of molecular layer interneurons. Plot shows the rate-corrected cross-correlogram13 denoting the relative firing rate of the first molecular layer interneuron aligned to the time of a spike in the second molecular layer interneuron at *t*=0 ms.

Across our complete database of cerebellar neurons, we found *n=23* examples where the simple spikes of ground-truth Purkinje cells show inhibition at monosynaptic latencies after a spike in a putative molecular layer interneuron (Figure 4C). The waveforms of the 23 putative molecular layer interneurons that inhibited a neighbor Purkinje cell (Figure 4D) form two groups: one group shows an early positivity with relatively little repolarization after the negativity; the other group shows initial negativity followed by a large positive deflection.

To identify putative molecular layer interneurons that may not directly inhibit Purkinje cells, we included all units with somatic spikes located in the molecular layer in our analysis. Consistent with prior reports^14,53^, we observe two classes of molecular layer interneuron waveforms (black versus gray) that could be separated by hierarchical clustering (Figure 4E). We found no evidence that molecular layer interneurons with a specific waveform shape were functionally connected to a Purkinje cell, with the caveat that we might have failed to document some molecular layer interneurons that inhibit Purkinje cells if we were not recording from a neighboring Purkinje cell at the same time.

Finally, in agreement with a previous report^53^, we found evidence for gap junction coupling in 2 of 16 simultaneously recorded pairs of molecular layer interneurons. The short latency peaks at −0.5 ms and +1.0 ms in the example cross-correlogram (Figure 4G) are indicative of gap junction coupling.

### Classification of granule layer elements

To provide expert labels for cerebellar units recorded in the granule cell layer, we began by considering action potential shape. Extracellular recordings from distal axons usually have thinner action potential waveforms than those recorded near the cell soma^54^. In our recordings from the granule cell layer, the distribution of peak-to-trough durations is strongly bimodal (D=0.06, p = 2.2 x 10^−16^) and many have thin action potentials with peak-to-trough durations shorter than 0.2 ms.

Within the waveforms from Figure 5A with durations shorter than 0.3 ms, many exhibited prominent negative-after-waves^20,55^. Our previous results with optogenetic activation of mossy fibers in mice^14^ established that the presence of a negative after-wave is sufficient, but not necessary, to identify mossy fibers *in vivo*. To be conservative, we used the presence of a negative after-wave (Figure 5B, arrow) as a necessary criterion for identification of putative mossy fibers. Many mossy fibers in our sample could fire at remarkably high firing rates, as indicated by the distribution of interspike intervals for an example unit (Figure 5C).

**Figure 5.**
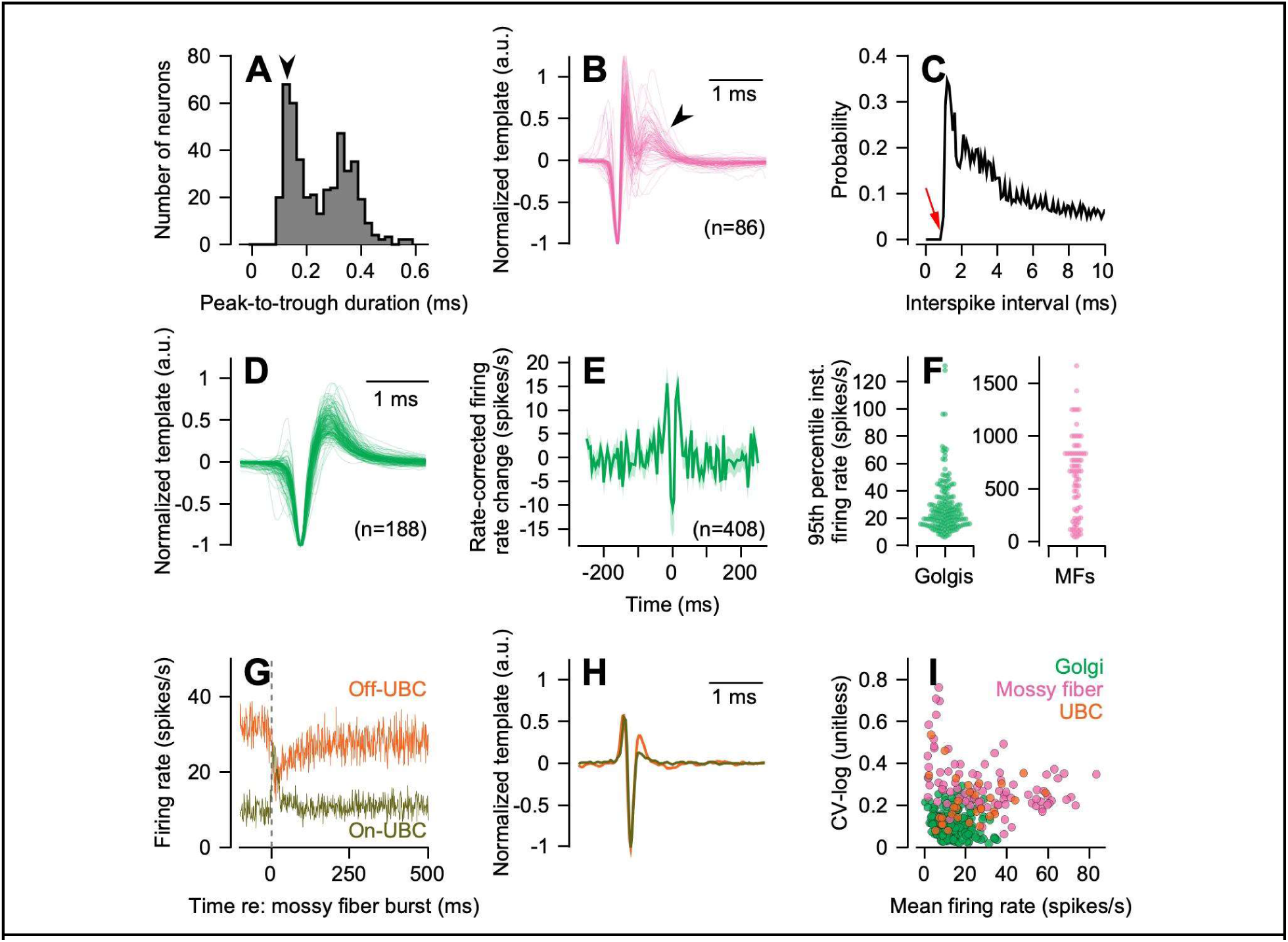
Characterization and identification of neural units in the granule cell layer. (A) Distribution of waveform durations on each neural unit’s primary channel, measured from the waveform’s peak to its trough. Arrow denotes the peak of the subpopulation of units with very brief waveforms. (B) Putative mossy fiber waveforms, with amplitude normalized. Arrowhead highlights the presence of a negative after wave. (C) Interspike-interval distribution for an exemplar mossy fiber. Red arrow highlights the short absolute refractory period of this fiber. (D) Normalized primary channel waveforms across a population of putative Golgi cells. (E) Rate-corrected cross-correlogram across unique pairs of simultaneously recorded Golgi cells. (F) Distribution of the 95th percentile of the instantaneous firing rate for putative Golgi cells (left, green) and mossy fibers (right, pink) shown as a swarm plot. (G) Exemplar putative unipolar brush cells (UBCs) identified functionally by their response aligned to bursts of simultaneously recorded mossy fibers. Olive and orange traces show an exemplar on- and off-UBC. (H) Primary channel waveforms for the On- and Off-UBCs shown in (G). (I). Scatter plot showing that CV-log and mean firing rate together do not discriminate granule layer neurons. Green, red, and orange symbols show data for putative Golgi cells, mossy fibers, and UBCs, defined by our criteria for expert identification.

A second distinctive class of action potentials recorded in the granule cell layer (Figure 5D) featured a stereotypically broad waveform, consistent with previously reported recordings from Golgi cells^20,56–60^. Their instantaneous firing rates provide additional evidence that the broad-waveform units in the granule cell layer are likely to be Golgi cells^56,57,60^ (Figure 5F). For each neuron, we measured the distribution of instantaneous firing rate, calculated from the inverse of each interspike interval across the full recording, and found the value of firing rate at the 95^th^ percentile of the distribution. This noise-immune measure of maximum firing rate was significantly different between putative Golgi cells and mossy fibers (independent samples t-test, t(244)=19.6, p = 4.7 x 10^−52^). Use of the 95th percentile as an estimate of maximum firing rate, rather than the mean or maximum instantaneous firing rate themselves, eliminates potential artifacts from the infrequent addition of spikes from noise or other neurons. Finally, across our sample of *n=188* putative Golgi cells, we saw limited evidence of gap-junction coupling in the form of millisecond synchrony of action potentials, previously reported *in vitro*^61^. However, simultaneously recorded pairs of Golgi cells (*n=408* pairs) showed some degree of synchronization over longer time scales (Figure 5E), potentially limited to longer time scales by active millisecond-scale desynchronization during behavior^62^.

A third group of neurons called unipolar brush cells (UBCs) exists in abundance in the vestibulocerebellum and portions of the cerebellar vermis^63,64^. To identify UBCs, we took advantage of known response properties from *in vitro* recordings^65^. Mossy fiber bursts driven by electrical stimulation *in vitro* cause post-synaptic responses in UBCs that span a range of time scales and can be depolarizing (on responses) or hyperpolarizing (off responses). Thus, we reasoned that we could identify putative UBCs in our sample of granule layer neurons by averaging the firing rates of units aligned on spontaneous, brief (<100 ms) bursts in mossy fibers that were recorded simultaneously. Putative on- and off-UBCs show responses with different time courses (Figure 5G) and have spike waveforms that are distinct from those of either mossy fibers or Golgi cells (Figure 5H).

The analysis in Figure 5 shows that action potential shape and functional properties such as the maximum instantaneous firing rate are likely to be quite informative about the identity of different neuron types recorded in the granule cell layer. In contrast, traditional approaches to cell-type identification in the cerebellum, such as plots of log-CV versus mean firing rate^33,34^, appeared unlikely to differentiate granule layer neuron types^66^ (Figure 5I). We noted earlier (Figure 2F) that a published classification method^32^ failed on our population of ground truth Purkinje cells; it assigned labels based on mean firing rate, local firing rate regularity assessed via CV2, and the median absolute deviation of interspike intervals from the median. Finally, unsupervised learning algorithms based on discrete waveform or firing metrics proved largely unsuccessful for disambiguating neuron types^66^, not a surprise given the unimodality of these features previously used to classify cerebellar cell types (Figure 1E). Therefore, we turn next to a classification strategy that was able to take advantage of multidimensional features.

### Potentially informative features of extracellular recordings

A deep-learning classifier, trained on a ground-truth library of neuron types determined in mice by optogenetic stimulation^14^, validated with >90% accuracy our expert labels for 585 units that were recorded in monkeys and classified according to the criteria outlined in Figures 2-5. Our next step is to ask which quantifiable features of our library of expert-classified neurons provide a basis for neuron-type identification. We consider measures of firing statistics, waveform, and local electrical effects.

#### Firing statistics

We developed an approach to assess firing statistics in a way that generalizes across tasks and species. We assess regularity properties independent of a neuron’s firing rate by constructing what we call “3D-ACGs”. We calculated the time-varying firing rate of each neuron and constructed separate ACGs, binned by firing rate decile, based on the local firing rate measured at each spike. For the example UBC shown in Figure 6, the 3D-ACG (Figure 6A) shows multiple bands of spike times that widen systematically as firing rate decreases, a pattern that is typical of a neuron with highly-regular firing rates that modulate reliably and strongly in relation to a behavior.

**Figure 6.**
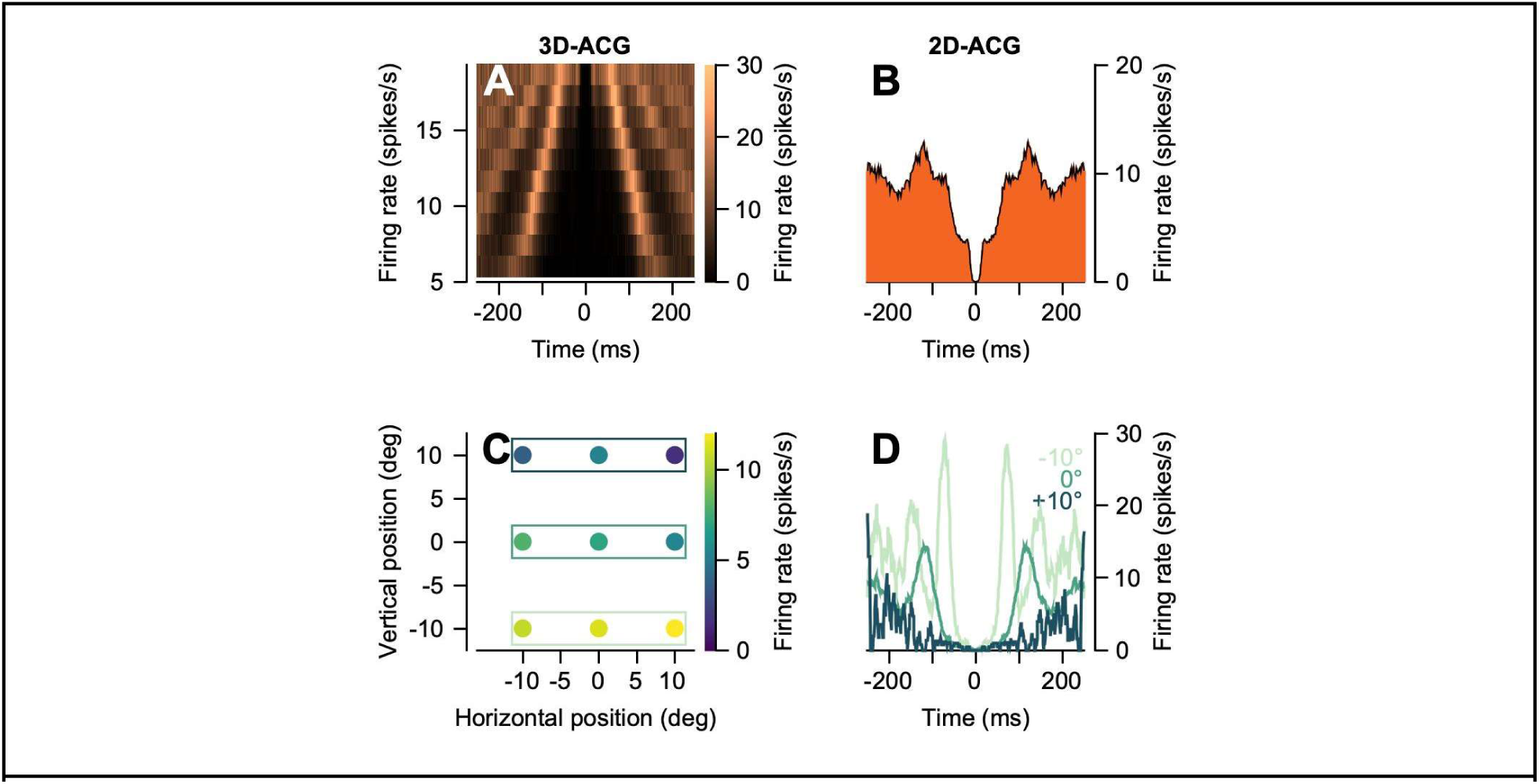
A tool to assess intrinsic regularity properties independent of stimulus- and response-related modulation of firing rate. (A) 3D-ACG for an example putative unipolar brush cell recorded in the granule cell layer. (B) Conventional (2D) auto-correlogram for the same neuron used in (A). The auto-correlogram is computed across the duration of the recording session. (C) The color axis shows the mean firing rate for the example UBC shown in A as the monkey fixated a stationary dot at each of nine points on a grid. (D) Conventional auto-correlograms for the same neuron shown in A-C, stratified based on the monkey’s vertical eye position. Colors of the traces in D correspond to the horizontal rectangles in C showing the monkey’s vertical fixation position.

The use of 3D-ACGs mitigates the impact of sensory stimuli and behavioral responses on scalar metrics of discharge regularity such as CV or CV2. As an example of the failures of the traditional ACG, we show analysis of a putative UBC recorded in the granule cell layer. The ACG computed across the duration of the experimental session (Figure 6B), without regard for the behavioral responses of the monkey, looks irregular and non-standard. The explanation is that firing rate varied systematically and strongly as the monkey varied its eye position. To demonstrate the relationship between firing rate and eye position, in a subset of trials the monkey fixated different stationary targets (Figure 6C). When the monkey fixated below the horizontal meridian (−10°), firing rate increased. Three ACGs contingent on the vertical fixation position (Figure 6C) have more traditional shapes but are quite different from each other. Not only the mean firing rate but also discharge regularity depended on the vertical position of the monkey’s eyes. When the monkey fixated below the horizontal meridian at −10° versus at 0°, the mean CV2 was 0.50 versus 0.68, corresponding to increased regularity for the targets located below the horizontal meridian.

Visual inspection of the 2D-ACGs (Figure 7A) separated according to their expert label showed reasonable consistency within neuron types and differences across neuron types. The same is true of 3D-ACGs (Figure 7B), though challenges of visualization preclude showing more than an example for each neuron type.

**Figure 7.**
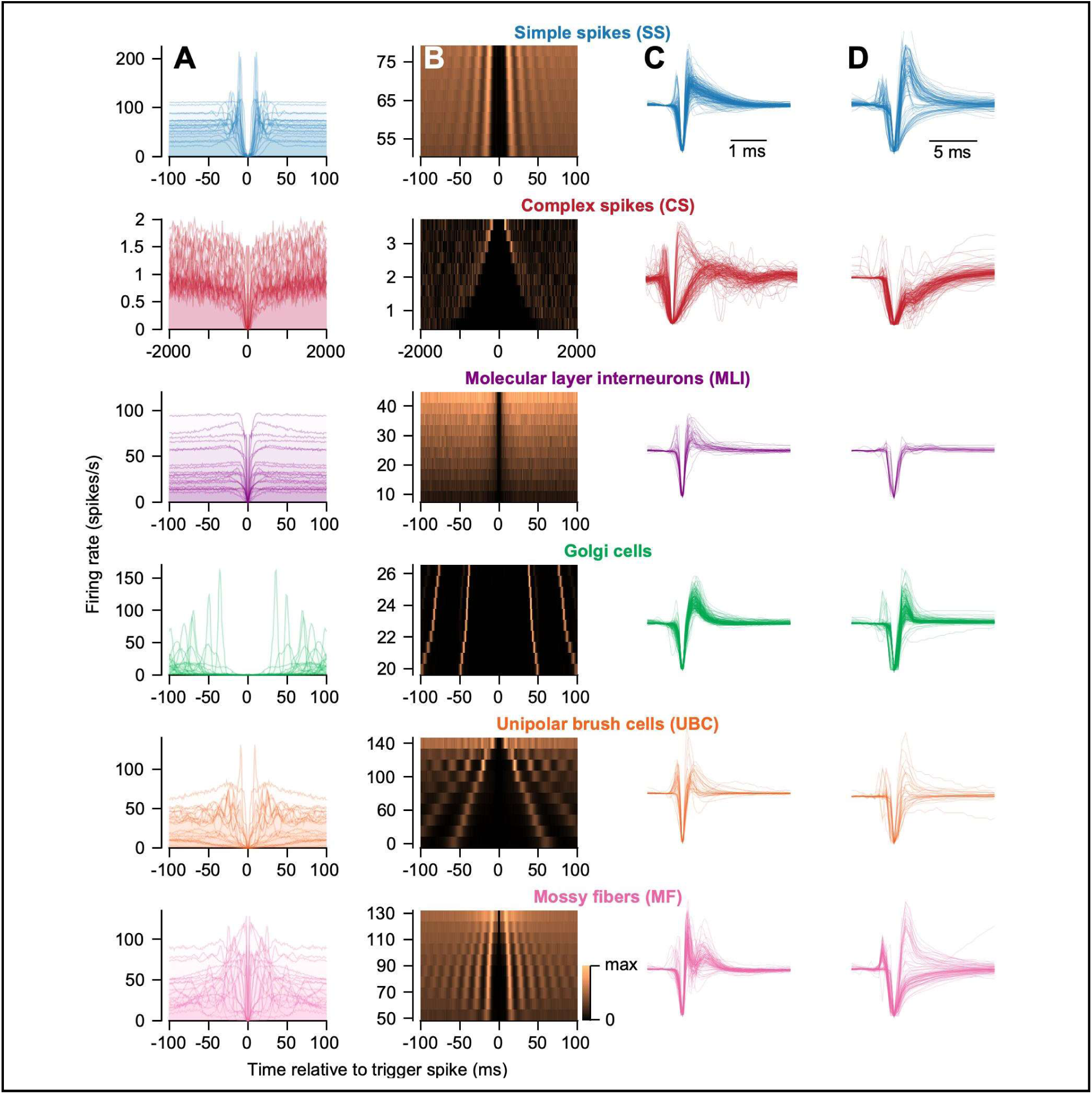
Features of expert-identified neurons in the primate cerebellum. (A) Conventional (2D) auto-correlograms for a random subset of *n*=40 neurons for each putative neuron type. (B) 3D-ACG for a representative example neuron of each cell type. (C) Primary channel waveform for all neurons of each type. (D) Spike-triggered LFP recorded on each neuron’s primary channel. Waveforms in (C) and spike-triggered LFPs in (D) have normalized amplitudes and potentially have been inverted, as described in the text.

#### Waveform

In our previous paper, we used the complete time series of a neuron’s primary channel waveform as feature for classification of ground-truth identified neurons in mice^14^, a feature that has proven useful for neuron identification across brain areas^38,40^. Visual inspection of Figure 7C and our previous unsupervised classification approach in Figure 3G suggests that waveform is likely to be a similarly useful feature for classification of neuron type in our expert-classified recordings.

#### Local electrical effects

We were inspired by previous attempts in the cerebellar literature to automatically identify complex spikes during spike sorting^67–69^ using the distinctive complex-spike triggered deviations of the LFP as one potential feature. To test the possibility that the LFP contains additional information about cell types not present in the typical spike band action potential, we quantified LFP deflections aligned to each neuron’s action potential. The resulting spike-triggered LFP time series was subsequently normalized and reflected, if necessary, using the same procedure we use for primary channel waveform. Inspection of Figure 7D again suggests strong similarity of the spike-triggered LFP within neuron types and systematic variation across neuron types. In evaluating Figures 7C and D, note the difference in the time scale of the traces in the two columns.

### Information about neuron type from different features of extracellular recordings

To achieve a quantitative answer to the question of which features of extracellular recordings are most informative about neuron type, we leveraged a deep learning classifier in combination with a principled approach for equalizing the dimensionality of different features. To mitigate differences in the overall parameter space for different inputs, we compressed each input into the same dimensionality latent vector using separate variational autoencoders^70,71^ (Figure 8A). We trained variational autoencoders to reconstruct each feature, one at a time, by sampling from a compressed 10-dimensional latent space (see Methods). Following training and optimization of the autoencoders, we could leverage a common classifier architecture to evaluate the information content of different features for cerebellar neuron type classification. Using leave-one-out cross validation, we quantified the information contained in the compressed space and asked whether each individual feature could correctly identify the type of the withheld neuron.

**Figure 8.**
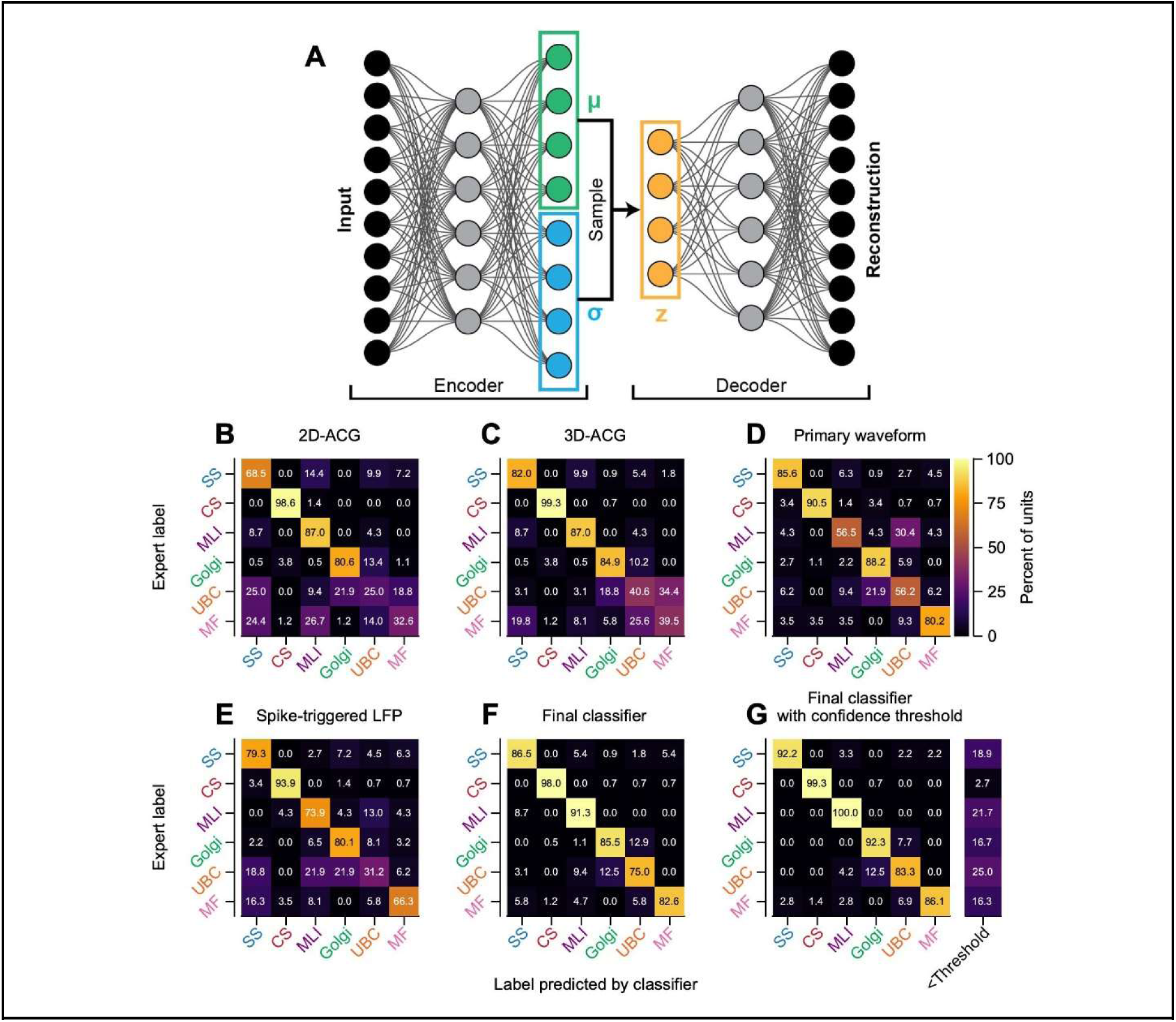
Assessment of classifier performance for expert-identified neurons across waveform and regularity features. (A) Deep-learning strategy for an unbiased quantification of information content for classification based on differently-sized input features. Diagram shows a variational autoencoder that encodes high dimensional inputs into a lower dimensional bottle neck (squared regions). A decoding arm learns to sample from the encoder and recapitulate the inputs. (B-E) Cross-validated classification performance for various extracellular features, each compressed via an optimized variational autoencoder. Each panel shows a “confusion matrix” where each column in the matrix reports, as percentages, the distribution of expert labels as a function of the neuron type predicted by the classifier. (F) Performance of a “full” classifier that takes 3 features as inputs: 3D-ACG, primary channel waveform, and spike-triggered LFP. (G) Full classifier performance when we threshold its output according to a confidence ratio14 computed across 25 training replicates. Far right column in (G) denotes the percentage of each expert-labeled neuron type that did not exceed the confidence threshold of 2.0.

Each of the features we tested was somewhat informative about neuron type. We quantify the performance of each feature with “confusion matrices” (Figure 8B-G) where each column in the matrix reports the distribution of expert labels as a function of the neuron type predicted by the classifier. We summarize each confusion matrix using the “micro-average” of assignments that agreed with expert classification across all 585 neurons in the sample. The use of micro-average accounts for different sizes of the samples by reporting the percentage of correct identification across the whole population, not the average across the diagonal. Of the features, the 2D-ACG (Figure 8B) was the least informative (micro-average: 73.0%), the 3D-ACG (Figure 8C) was more informative (micro-average: 79.0%), the waveform (Figure 8D) was most informative (micro-average: 84.1%), and spike-triggered LFP (Figure 8E) was comparable to 3D-ACGs (micro-average: 78.5%).

More granular examination revealed that different metrics were less informative for different neuron types, lending hope that they would be more informative together. Using either 2D-ACG or 3D-ACG alone, the classifiers performed poorly on mossy fibers and unipolar brush cells. Using only waveform, the classifier performs better compared to 2D-ACG or 3D-ACG at identifying mossy fibers. However, there was significant classifier confusion between molecular layer interneurons and UBCs using waveform. In contrast, classification using spike-triggered LFP (Figure 8E) showed less confusion between molecular layer interneurons and UBCs (13% versus 30.4% misclassified).

Given that different features appear to contain complementary information about neuron-type (Figure 8B-E), we tested whether classification on the combination of all inputs would achieve robust performance. Indeed, a classifier that took three features (3D-ACG, primary channel waveform, and spike-triggered LFP) as inputs resulted in an improvement to a micro-average classification performance of 88.0% across all neurons (Figure 8F). Classifier performance was improved further to 93.2% (Figure 8G) by applying a threshold on the relative confidence of classifier labels^14^. Application of a confidence threshold caused fewer than 20% of the units we recorded to be excluded from our labeled sample. Overall, we conclude that single metrics, even if high dimensional, are helpful but insufficient to obtain accurate neuron-type identification. Rather, the multiple metrics available to us appear to encode complementary information and together they allow automated classification of neuron type in cerebellar recordings.

## Discussion

Understanding the processing in neural circuits requires the ability to identify the information transmitted between neuron types^4,8,12^. Here, our goal was to use a combination of logic, circuit architecture^3,18^, and prior observations^19,20,35,55,58–60^ to assign labels to cerebellar neurons recorded in behaving primates. Further, we provide a quantitative analysis of the information about neuron type in various readily-accessible features of extracellular recordings in the cerebellum. Together, these steps form the foundation for understanding circuit-level processing in the service of complex cerebellar-dependent behaviors.

Our approach succeeded. A cascade of objective criteria allowed automated identification of 6 cerebellar neuron-types from their extracellular features: Purkinje cells, climbing fibers, molecular layer interneurons, Golgi cells, mossy fibers, and unipolar brush cells. While we did not obtain ground-truth identification through optogenetics, our expert neuron-type identification agreed impressively with ground-truth identification in mice^14^. Further, we used machine-learning technology^70,71^ to verify that the units we identified by expert criteria clustered on the basis of electrophysiological features, and that those features were quite informative about neuron type. Thus, we are quite confident in our expert neuron-type identification and we now possess an automated approach to identify neurons that were not tested with the explicit criteria used for our original expert-identification. Next steps are: (i) use knowledge of neuron-type to evaluate how the cerebellar circuit computes and learns; (ii) extend cerebellar neuron-type identification beyond the neuron types we already can classify; (iii) facilitate application of the same neuron-type identification strategy to non-cerebellar structures with similar richness of neuron types.

### A strategy for expert identification of cerebellar neuron type

We are confident that our conservative approach ensures that we assigned the correct label to the almost all of our neural units. Our strategy to assign an expert label to an individual neural unit required a preponderance of evidence. We used different criteria to identify different neuron types, drawing from the layer of the recording, firing rate statistics, functional interactions with other identified units, and waveform shape. Regardless, label assignments for all neuron types (beyond ground-truth Purkinje cells and complex spikes) required some degree of subjective assessment of the available information.

We took pains to ensure that our sample of neurons represented well-isolated single units because the criteria we used to disambiguate neuron types relied on features of extracellular recordings, such as primary channel waveform and regularity properties. During post-sorting curation of the recordings, we removed any units with low signal-to-noise ratios, evidence of instability across the recording session, refractory period violations, or any other evidence of contamination by multi-unit activity. Differences in our ability to obtain sufficiently isolated and stable recordings likely biased the number of units in our sample across neuron classes. For instance, the relatively small sample sizes of molecular layer interneurons and UBCs (*n=32*) might be due to their smaller size relative to other neurons in the cerebellar circuit.

We used functional interactions within the cerebellar circuit to identify several neuron types. For instance, we identified a class of putative molecular layer interneurons by their interaction with ground-truth Purkinje cells, as assayed via their cross-correlogram. A statistically-significant, properly-timed inhibition of Purkinje cell simple spikes seems like a definitive metric to identify molecular layer interneurons. Yet, the stringent criteria of monosynaptic inhibition of a simultaneously recorded Purkinje cell likely excludes many molecular layer interneurons because either a Purkinje cell was not recorded simultaneously, the recorded Purkinje cell was not a target for the molecular layer interneuron under study, or the molecular layer interneuron might solely inhibit other molecular interneurons rather than Purkinje cells^53^. Therefore, while monosynaptic inhibition of a Purkinje cell is likely sufficient for identification of molecular layer interneurons, we also used the layer of the recording to establish the identity of molecular layer interneurons in the absence of such functional interactions.

To assign units as putative mossy fibers, we required the presence of a negative after wave on the neuron’s primary channel waveform, corresponding to a recording near a glomerulus^14,55^. As a negative after wave was not always present in ground-truth mossy fiber recordings^14^, we assume that we excluded from our sample a subset of mossy fibers not recorded near the glomerulus.

### Automated identification of cerebellar neuron-type

Our choice to move forward with new approaches to separate cerebellar neuron classes was reinforced by 1) the poor classification performance of the previous algorithm based on data from awake and anesthetized rabbits^32^, 2) the unimodal nature of the discrete metrics previously used for cerebellar classification in awake monkeys^33,34^, and 3) our failure to derive meaningful clusters using unsupervised techniques.

The challenge we set out to address is to provide accurate and reliable identification of neuron type using approaches that do not depend on subjective judgements by self-acclaimed experts. Validation by the ground-truth classifier developed for mice^14^ implies that our conservative reliance on the preponderance of evidence yielded accurate neuron-type identification. Even across species and distinct cerebellar areas, classifiers trained on ground-truth identified neurons in mice predicted labels that agreed on more than 90% of neurons with our “expert” labels.

Armed with a believable set of identities for our expert-classified sample, our next step was to deploy deep learning to ask which features of electrophysiological recordings are informative about neuron-type. We then developed a classifier that can be used for automated neuron-type identification in larger samples of neurons.

Overall, we found that different features derived from extracellular recordings provide complementary information about neuron-type. All the electrophysiological features we tested were quite informative, but their weaknesses appeared for different types of neurons:

- 3D-ACGs provided comparable or improved classification performance across all classes compared to traditional ACGs: yet both performed poorly on mossy fibers and unipolar brush cells. As a tool for neuron-type classification, it is likely that the superiority of 3D-ACGs is a general finding. At the very least, in the case where a neuron’s activity is largely unmodulated across a recording, the resulting 3D-ACG would have the same information as a traditional ACG.
- Spike-triggered LFP was able to distinguish molecular layer interneurons and UBCs while waveform was particularly useful for identification of mossy fibers. It is likely that waveform and spike-triggered LFP represent different ‘views’ of the same neuron because of their different, albeit partially overlapping, frequency content. The spike-triggered LFP depends on a combination of neuron morphology^72^ and post-synaptic/circuit-level effects^47,73^. Therefore, it seems reasonable that LFP signals would differ across neuron types given their different locations in the cerebellar connectome^3,18^.

Not surprisingly, a combination of all features was more informative about neuron type than any of the individual features. Further, our final classifier performed at greater than 90% accuracy on the expert-classified dataset, especially when we insisted on a threshold for classifier confidence. We conclude that we can use the combined classifier in the future to identify neuron type.

### Cerebellar layer identification

Identification of the layer of a recording is extremely useful for neuron-type identification. Here, we used two tools that have proven useful for layer identification in the cerebral cortex and demonstrated that both current source density analysis^44–50^ and the normalized LFP^52^ allow identification of cerebellar layers. The current source density analysis requires a behavior or sensory stimulus that drives activity to temporally align individual trials^44,52^; a challenge for some studies. The normalized LFP^52^ removes the necessity to identify a temporally discrete modulatory sensory or behavioral stimulus, but introduces other limitations. For instance, the bands of activity are less distinct and layers more difficult to identify in an electrode penetration that crosses multiple layer boundaries. With short electrodes, recordings that do not span cerebellar layers due to the orientation of the electrode relative to the laminar structure of the cerebellum will be challenging to interpret. Recording with more contacts or longer probes might exacerbate the problem of multiple layer crossing, though the use of “local” rather than “global” normalization might mitigate some of these issues. Finally, we note that both the current source density and normalized LFP analyses require measurements of differential LFP activity across recording contacts; neither analysis is possible with recordings from single electrodes.

### Known unknowns in the cerebellar circuit

Several cerebellar neuron types either are inaccessible to extracellular recordings or are sufficiently rare in number that we don’t seem to have recorded a sufficient sample. For instance, we don’t think we can record granule cells on our current probes given the relatively low impedance (1-2 MΩ) and contact size (7.5 µm diameter) of our electrodes, as well as the small size, closed electrical field, and high density of granule cells. Further, relatively rare cerebellar cell types, such as Purkinje layer interneurons^18,74^ and candelabrum cells^75,76^, escaped our ability to identify and label them. The relative dearth of information about the connectivity profiles, electrical signatures, and response properties of these neuron types makes assignment of expert labels to them impossible at this time.

### Applicability to other brain regions

Can the methods and procedures outlined here to identify cerebellar neurons also be applied to other regions of the brain? We believe that our strategy is general enough to potentially disambiguate cells in other brain regions. For instance, waveform shape contains information for neuron type identification in the cerebral cortex^38,40,77–80^. Differences in action potential shape and regularity properties are inherently the result of differences in morphology, ion channel content, and circuit connectivity. We think that spike-triggered LFP might be particularly informative in non-cerebellar structures, including those without clear layers. Therefore, in brain regions where neuron types of interest show distinct anatomical, connectivity, or ion channel dynamics, the methods we outline here may be sufficient to “label” and subsequently classify neuron type.

## Acknowledgements

Our research is supported by NIH grants R01-NS112917 (SGL) and K99-EY030528 (DJH). We thank Stefanie Tokiyama and Bonnie Bowell for monkey assistance. We are grateful to members of the Cerebellar Cell-type Classification Collaboration (C4) for helpful comments and discussions.

## Author contributions

DJH and SGL designed all experimental procedures. DJH performed recordings in the cerebellar flocculus. DJH analyzed the data. NJH developed the spike-sorter and collaborated on data analysis and visualization. DJH and SGL designed the figures and wrote the manuscript.

## Methods

All experiments were performed on three rhesus macaques (*macaca mulatta*, male, 10-15 kg). A portion of the dataset described in this study was reported in two previous publications^13,14^. All experimental procedures were approved in advance by the Duke *Institutional Care and Use Committee* (Protocols A085-18-04, A062-21-03, and A016-24-01) and performed in accordance with the *Guide for the Care and Use of Laboratory Monkeys* (1997).

### General procedures

Each monkey underwent several surgical procedures prior to data acquisition. Each surgical procedure was performed using sterile technique while the monkey was deeply anesthetized with isoflurane. Monkeys received analgesics post-op until they had recovered. In the first surgical procedure, we implanted a head-restraint system that would allow us to measure eye movements uncontaminated by changes in head position. In a separate surgery, we sutured a small coil of wire to the sclera of one eye^81^, allowing us to measure the monkey’s eye position with high temporal and spatial precision using the search coil technique^82^. The monkey subsequently was trained to perform discrete trials of smooth pursuit eye movements in exchange for a fluid reward. Once the monkey had demonstrated proficiency in tracking the visual target with minimal intervening saccadic eye movements, we performed a final surgical procedure to implant a recording cylinder allowing electrode access to the floccular complex of the cerebellar cortex. We implanted the recording cylinder 11 mm lateral to the midline, angled 26° backwards from the coronal plane, and directed at the interaural axis.

### Behavioral procedures

The general procedures for recording monkeys’ smooth pursuit behavior have been described in detail previously^13^. Briefly, monkeys were seated in a dimly lit room with their heads fixed 30 cm in front of the CRT monitor (2304×1440 pixels with an 80 Hz refresh rate). Visual targets (0.5° diameter black spots) were presented on the monitor in discrete trials, controlled by our lab’s custom Maestro software. During some ‘fixation-only’ trials, the visual target appeared in one of nine discrete locations (spanning a 10°x10° visual square). The monkey received a small liquid reward for fixating the target within ±1° for one second. The vast majority of the experimental session consisted of discrete trials of smooth target motion. The target appeared in the center of the screen at the start of each trial. The monkey was required to maintain fixation on the target within an invisible bounding box of ±3° for a uniformly random interval of 400 to 800 ms. At the end of the fixation interval, we shifted the position of the target in one direction by 0.15|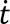| degrees and moved it smoothly in the opposite direction at a constant velocity of 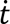 degrees/sec^83^. The backwards step minimizes the number of catch-up saccades during the initiation of the smooth pursuit eye movement^84^. All monkeys had extensive experience performing smooth pursuit tasks prior to data collection. With the exception of the current source density analysis (see below), analyses were not contingent on the task-related performance of the monkeys. We digitized separately at 1 kHz the horizontal and vertical position of the monkey’s eyes as measured from the scleral coil system and stored the data for later offline processing.

### Neurophysiology procedures

We acutely inserted either tungsten micro-electrodes (FHC, ∼1 MΩ) or, more commonly, custom manufactured Plexon S-Probes through a craniotomy into the floccular complex of the cerebellum. Plexon S-Probes had 16 contacts arranged in two columns on a grid with spacing at 50 µm. Each contact was a tungsten micro-wire with a diameter of 7.5 µm. Each day, we drove the electrode through the cerebellar cortex using a Narishige microdrive (MO-95/MO-97) with the goal of recording activity from the region of the flocculus and ventral paraflocculus that controls smooth eye movement, a region that we call the floccular complex. We recognized the floccular complex by its strong response to smooth pursuit eye movements as well as the occasional occurrence of Purkinje cell complex spikes. After arriving in the floccular complex, we waited a minimum of 30 minutes (up to several hours) before recording extracellular spiking activity. The waiting period maximized the signal-to-noise ratio and minimized the drift of neural units across the electrode during the recording.

We used a 4-pole low-pass Butterworth hardware filter prior to digitization of continuous voltage signals from the contacts of the recording electrode to ensure that the voltage signals were uncontaminated by interference from the scleral coils. Wideband data were digitized continuously at 40 kHz using the Plexon Omniplex system.

After each recording session, we post-processed the data by applying a 300 Hz high-pass first-order Butterworth filter to the continuous wideband data recorded on each electrode. This preprocessing step mimicked the hardware filter used on Neuropixels probes, allowing comparisons between the neurophysiological signatures of our data in the monkey with our previously reported results across species^14^. Following pre-processing, we assigned individual action potentials to neural units using the semi-automated “Full Binary Pursuit” (FBP) spike-sorter^85^. As we were interested in leveraging potential monosynaptic interactions between simultaneously recorded neural units as a criterion for expert labeling, we chose FBP due to its superior ability to disambiguate action potentials that are within close temporal and spatial proximity. Following sorting, we manually curated the sorted units to ensure that each had a high signal-to-noise ratio and a low percentage of interspike interval contamination. We defined the signal-to-noise ratio based on the peak-to-trough amplitude of the waveform on the primary channel relative to the standard deviation of the noise on that channel, computed as 1.96 times the median absolute deviation of the complete voltage timeseries on the primary channel. Use of the median absolute deviation to compute the standard deviation of the channel noise ensured that voltage fluctuations due to action potentials did not bias our estimate of the noise amplitude. We defined the percentage of ISI violations by determining the percentage of spikes that occurred during an assumed absolute refractory period of 1 ms. The 1 ms assumed refractory period represents an upper bound on the percentage of ISI violations as we were able to routinely isolate putative mossy fibers whose instantaneous firing rates intermittently exceeded 1,000 spikes/second.

We computed the LFP time series by applying a causal 2nd order bandpass Butterworth filter to the wideband voltage recordings (high-pass cut-off: 5 Hz, low-pass cut-off: 500 Hz). We subsequently downsampled the filtered voltage time series to 2500 Hz. We chose these parameters to mimic the parameters of Neuropixels recordings, although we note that we implemented an additional high-pass filter to minimize interference from very low frequency signals.

#### Auto- and cross-correlograms

We computed conventional auto- and cross-correlograms in the same manner as we described previously^13^. Briefly, we computed the probability of observing a spike in millisecond-wide bins relative to a ‘trigger spike’. For an auto-correlogram, we considered each spike as the trigger spike and then measured the probability of the same neuron spiking at each millisecond relative to that spike. We normalized the probability by the bin size (1 ms, 1000x) to ensure that the shape and magnitude of the auto-correlogram were independent of the chosen bin size and to convert the units of the auto-correlogram to spikes/second. By convention, we set the *t=0* ms bin to zero when computing auto-correlograms. We computed cross-correlograms in the same manner, except we assayed the probability of spiking in a second neuron, N2, relative to the time of each spike in a first neuron, N1:

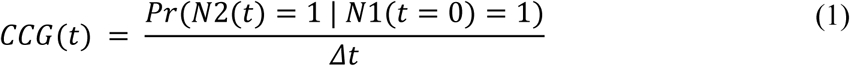

In Equation 1, the probability of N2 firing rate some time *t* is assessed relative to each spike of N1. The bin width, Δ*t* (1 ms), in the denominator expresses the CCG in units of spikes/second.

#### 3D auto-correlograms

Our goal was to identify the intrinsic regularity properties of units without contamination by stimulus-related or movement-related changes. Our approach centers on the construction of stacked (3D) auto-correlograms that are stratified by the local firing rate responses of each neuron spike. We described the general process to construct a 3D-ACG previously^14^. Briefly, we computed the instantaneous firing rate of each neuron across the complete recording session using the inverse interspike interval method^86^. We then smoothed the instantaneous interspike interval using a noncausal boxcar filter with a width of 250 ms. We measured the value of the smoothed instantaneous firing rate time series at the time of each action potential. The resulting distribution of smoothed instantaneous firing rates were divided into equal sized deciles. We computed separate conventional auto-correlograms for each decile by selecting the spikes used as the trigger spike (i.e., *t=0* ms) whose smoothed instantaneous firing at the time of the trigger spike fell in each decile. We visualized the 10 resulting auto-correlograms as a surface, where the color axis corresponds to the firing rate computed from individual ACGs via Equation 1, the x-axis corresponds to the time relative to the trigger spike, and the y-axis corresponds to the firing rate decile from the slowest firing rate to the fastest.

### Classification of neuron type

Previous studies largely focused on a combination of scalar metrics to disambiguate neuron types both in the cerebellum^32–34^ and in other areas of the brain^36,80^. While such metrics have proven successful in some instances, they are often not robust^40,66^ to different recording methodologies, laboratory procedures, or species. Therefore, our approach^14^ is to leverage semi-raw data to establish robust, albeit high-dimensional, metrics for neuron-type classification.

#### Neuron waveforms and spike-triggered local field potentials

Mean waveforms were computed following spike sorting by applying the drift-shift algorithm^14,87^ to correct misalignments in spike sorter output on a spike-by-spike basis and avoid adverse effects of spike timing jitter on the mean waveform. The drift-shift algorithm also strategically chooses individual action potentials to average for the mean waveform, with the goal of removing very low (potential noise) or very high amplitude (potential artifacts) events from the mean waveform.

Briefly, we specified the primary channel of each spike as that with the largest peak-to-trough amplitude. We selected up to 5,000 individual spike events whose primary channel corresponded to the unit’s overall primary channel, using only events up to the 95% percentile of spike amplitudes to avoid potential inclusion of high amplitude artifacts in the average. Then, we iteratively shifted individual action potentials to maximize the cross-correlation across the sample of action potentials. Finally, we used the mean across the selected and time-shifted action potentials as the neuron’s mean waveform. We note that our results do not depend on the drift-shift algorithm. Because our sample contained mainly units with high signal-to-noise ratios and minimal drift across contacts, the drift-shift aligned waveforms appeared qualitatively similar to those obtained by simply averaging the output from spike-sorting. We used a similar procedure to measure the mean spike-triggered LFP response. Here, we downsampled the spike times of each unit as measured by the spike sorter to 2500 Hz, corresponding to the sampling rate of our LFP. We used the primary channel for spikes as the primary channel for the LFP and otherwise aligned individual LFP “clips” in the same manner as traditional spikes.

For all subsequent analyses of waveform and spike-triggered LFP, we normalized the amplitude of the voltage trace. Normalization is important for both visualization as well as classification, as amplitude differences are due primarily to proximity of the recording contact to the neuron. If necessary, we inverted the neuron’s mean waveform/LFP to ensure that the primary deflection used for normalization was negative.

#### Current source density and local field potential analysis

We performed current source density or normalized LFP analysis only for recordings made with 16-contact S-Probe recordings, as those analysis techniques are not amenable to single-channel recordings. For the 16-contact recordings, we began by averaging the LFP time series across the two columns of contacts, yielding eight LFP time series, one for each row of contacts (spacing 50 µm). We filtered each contact’s LFP signal in time using a 3rd-order Savitzky-Golay filter and subsequently computed the current source density as the second spatial derivative of LFP signal across contacts using a 2nd-order Savitzky-Golay filter. We temporally aligned the resulting derivative map to the onset of target motion during discrete smooth pursuit trials.

Alignment was not contingent on the direction of target motion, but all visual stimuli moved at a constant speed of 20 °/sec. For the purpose of visualization, we upsampled the measured current source density at 5 µm resolution using 2D-spline interpolation.

We computed the normalized LFP (the “spectrolaminar pattern”) using established procedures from the macaque cerebral cortex^52^. Briefly, after averaging the LFP signal across columns, we computed the spectral power at each frequency (resolution 2.5 Hz) using the multi-taper method^88^. We smoothed the resulting power estimate in the frequency domain using a boxcar filter with a 25 Hz width. Finally, for each frequency bin, we computed the normalized LFP response by dividing by the maximum power across channels according to Equation 2:

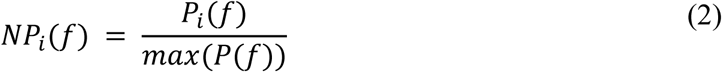

In Equation 2, *NP_i_*(*f*) represents the normalized power of the *i*-th contact in the *f*-th frequency bin. The normalized power on each channel was computed as the measured power of that frequency on the *i*-th contact, *P_i_*(*f*), divided by the maximum power in that frequency bin measured across all contacts.

#### Assaying information for classification using variational autoencoders

Our goal was to quantitatively measure the information present in high dimensional features that could be used for classification of cerebellar neuron types. Yet, the features that we wished to measure had different dimensions and different modes of information content. For instance, primary channel waveform in our dataset was represented by a single time series (160 elements) whereas a 3D-ACG was represented by an image with much higher dimensionality (10 x 250 pixels). To quantify the information content present in these various inputs, we devised a strategy to perform principled dimensionality reduction and compress the input feature space into a lower dimensional representation. A common-sized low dimensionality representation of each input space then could be used directly in a simplified classification architecture with a structure that was chosen *a priori*. Together, the common input space and shared classification architecture equalized the number of fitted parameters across classification models and ensured that we were not overfitting the classifier on our dataset. Thus, the common classification framework allows direct comparison of each low dimensional feature space to classify cerebellar neuron cell types.

We used variational autoencoders to reduce the unconstrained size of each input parameter into a lower dimensional (10-element vector) representation^71,89^. We reasoned that the demixing nature of the variational autoencoder would result in improved classification performance compared to traditional autoencoders. We trained a separate variational autoencoder for each type of high dimensional feature (waveform, spike-triggered LFP, auto-correlogram, and 3D-ACG). For each autoencoder, we used our full sample of neurons recorded from the floccular complex (*n=1,152*), including both neural units that had an expert label (*n=585*) as those that did not have an assigned expert label (*n=567*). The autoencoder was trained via stochastic gradient descent^90^ to minimize a cost function that included the weighted contribution of the mean squared error of the reconstruction from the input as well as the deviation of the low dimensional representation from a set of standard Gaussians (zero mean, unit variance) using the Kullback-Leiber divergence^71^.

The relative weights of these two error terms were set using β-normalization^70^ and modulated using a cosine annealing schedule^91^ that improves convergence during training. The total cost function corresponds to the evidence lower bound (ELBO), which was minimized across iterations. For each input type, we hand-tuned both the autoencoder architecture and parameters (e.g., number and size of hidden layers, learning rate, size of convolutions, type and size of pooling layers) to minimize the total cost as assayed on a withheld validation sample consisting of 30% of the complete sample of recorded neurons. This optimization procedure ensured that we had maximized the amount of compressed information in the low-dimensional encoded representation while simultaneously ensuring demixing of the low-dimensional representation.

After optimizing the variational autoencoder architecture for each respective high-dimensional input type, our goal was to quantify the amount of information in the compressed representation that could be used to classify different cerebellar cell types. As each representation was 10-dimensional, we could use identical architectures and training procedures to evaluate classification performance across features. The architecture and training procedure for our classifier was established *a priori* to ensure an unbiased comparison across inputs. The classifier used a multi-layer perceptron network, consisting of a 10-dimensional input layer (to receive the output of the optimized variational autoencoder latent representation), a 100-unit hidden layer with rectified linear activation functions^92^, and an ultimate output layer with a softmax activation function. Each element in the output layer corresponded to a single expert-identified cerebellar cell type.

We evaluate classifier performance using leave-one-out cross validation. For each neuron, we trained 25 models with random initial conditions. We split the remaining *n-1* expert-labeled neurons into separate training (70%) and validation (30%) sets. Because our expert-labeled dataset contained an unequal number of samples in each neuron class, we randomly downsampled over-represented classes to ensure they represented no more than 2-fold the number of samples in the smallest class. We then used random over-sampling to resample any under-represented classes, thereby equalizing the number of samples per class. Finally, we trained our multi-layer perceptron classifier using stochastic gradient descent^90^ to minimize the cross-entropy computed on the validation set. We used early termination to stop the training procedure when the cross-entropy as evaluated on the withheld validation set increased for more than five iterations. This “early stopping” procedure was implemented to prevent over-fitting to the training set and thereby promote generalization. Following training, we evaluate the prediction for the left-out neuron for each of the 25 random replicates of the classifier model.

After we had established the complementary information content across available input types, we trained a classifier that used multiple inputs to optimally classify our expert-labeled cerebellar cell types. As above, we used leave-one-out cross-validation to evaluate the performance of our ultimate classifier. For each withheld neuron, we trained a multi-armed neural network to predict the cell type labels of the remaining neurons. One arm of the neural network featured a convolution neural network whose architecture was identical to the penultimate latent layer of optimal convolution neural network we established by training the 3D-ACG variational autoencoder. The second and third arms provided inputs for the normalized waveform and spike-triggered LFP. Each arm supplied input to a common 100-unit hidden layer with rectified linear activation functions. Again, the final layer of the merged classifier featured a single unit per cell type with a softmax activation function. Training with early stopping proceeded as above and was terminated when the cross entropy of the validation set increased for consecutive training iterations. We repeated this procedure 25 times for each with-held neuron, providing an ensemble of models^93^ with different initial conditions and training and validation sets.

To threshold the output of our final classifier based on the ‘confidence ratio’, we used a previously established technique^14^. For each of the 25 randomly instantiated classifier models per leave-one-out sample, we obtained separate probability distributions for each cell type by aggregating the softmax outputs of each classifier. Dividing the mean of the most probable cell type distribution by the mean of second most probable cell type provided us with the confidence ratio. Neurons with a confidence ratio less than 2, indicating that two cell-type labels had similar mean probabilities, were deemed unclassifiable (below threshold) and thus were not included our evaluation of classifier accuracy.

## Data availability

All data for this study have been deposited into the Open Science Framework Database by the date of publication. Additional requests for data can be made to the corresponding author.

## References

1. Lisberger, S. G. Visual Guidance of Smooth Pursuit Eye Movements. Annu. Rev. Vis. Sci. 1, 447–468 (2015).

2. Lisberger, S. G. Visual Guidance of Smooth-Pursuit Eye Movements: Sensation, Action, and What Happens in Between. Neuron 66, 477–491 (2010).

3. Ramón y Cajal, S. Histologie Du Système Nerveux de l’homme & Des Vertébrés. (Maloine, Paris, 1909). doi:10.5962/bhl.title.48637.

4. Luo, L., Callaway, E. M. & Svoboda, K. Genetic Dissection of Neural Circuits. Neuron 57, 634–660 (2008).

5. Fishell, G. & Heintz, N. The Neuron Identity Problem: Form Meets Function. Neuron 80, 602–612 (2013).

6. Ecker, J. R. et al. The BRAIN Initiative Cell Census Consortium: Lessons Learned toward Generating a Comprehensive Brain Cell Atlas. Neuron 96, 542–557 (2017).

7. Zeng, H. & Sanes, J. R. Neuronal cell-type classification: challenges, opportunities and the path forward. Nat. Rev. Neurosci. 18, 530–546 (2017).

8. Migliore, M. & Shepherd, G. M. An integrated approach to classifying neuronal phenotypes. Nat. Rev. Neurosci. 6, 810–818 (2005).

9. Gouwens, N. W. et al. Classification of electrophysiological and morphological neuron types in the mouse visual cortex. Nat. Neurosci. 22, 1182–1195 (2019).

10. Poulin, J.-F., Tasic, B., Hjerling-Leffler, J., Trimarchi, J. M. & Awatramani, R. Disentangling neural cell diversity using single-cell transcriptomics. Nat. Neurosci. 19, 1131–1141 (2016).

11. Josh Huang, Z. & Zeng, H. Genetic Approaches to Neural Circuits in the Mouse. Annu. Rev. Neurosci. 36, 183–215 (2013).

12. Masland, R. H. Neuronal cell types. Curr. Biol. 14, R497–R500 (2004).

13. Herzfeld, D. J., Joshua, M. & Lisberger, S. G. Rate versus synchrony codes for cerebellar control of motor behavior. Neuron 111, 2448–2460.e6 (2023).

14. Beau, M. et al. A deep-learning strategy to identify cell types across species from high-density extracellular recordings. 2024.01.30.577845 Preprint at 10.1101/2024.01.30.577845 (2024).

15. Marchal, G. A. et al. Recent advances and current limitations of available technology to optically manipulate and observe cardiac electrophysiology. Pflüg. Arch. - Eur. J. Physiol. 475, 1357–1366 (2023).

16. Mermet-Joret, N. et al. Dual-color optical activation and suppression of neurons with high temporal precision. eLife 12, (2023).

17. Tremblay, S. et al. An Open Resource for Non-human Primate Optogenetics. Neuron 108, 1075–1090.e6 (2020).

18. Hull, C. & Regehr, W. G. The Cerebellar Cortex. Annu. Rev. Neurosci. 45, 151–175 (2022).

19. Lisberger, S. G. & Fuchs, A. F. Role of primate flocculus during rapid behavioral modification of vestibuloocular reflex. II. Mossy fiber firing patterns during horizontal head rotation and eye movement. J. Neurophysiol. 41, 764–777 (1978).

20. Prsa, M., Dash, S., Catz, N., Dicke, P. W. & Thier, P. Characteristics of Responses of Golgi Cells and Mossy Fibers to Eye Saccades and Saccadic Adaptation Recorded from the Posterior Vermis of the Cerebellum. J. Neurosci. 29, 250–262 (2009).

21. Kase, M., Miller, D. C. & Noda, H. Discharges of Purkinje cells and mossy fibres in the cerebellar vermis of the monkey during saccadic eye movements and fixation. J. Physiol. 300, 539–555 (1980).

22. Noda, H. Visual Mossy Fiber Inputs to the Flocculus of the Monkey*. Ann. N. Y. Acad. Sci. 374, 465–475 (1981).

23. Noda, H. Mossy fibres sending retinal-slip, eye, and head velocity signals to the flocculus of the monkey. J. Physiol. 379, 39–60 (1986).

24. Ohtsuka, K. & Noda, H. Burst discharges of mossy fibers in the oculomotor vermis of macaque monkeys during saccadic eye movements. Neurosci. Res. 15, 102–114 (1992).

25. Gilbert, P. F. C. & Thach, W. T. Purkinje cell activity during motor learning. Brain Res. 128, 309–328 (1977).

26. Stone, L. S. & Lisberger, S. G. Detection of tracking errors by visual climbing fiber inputs to monkey cerebellar flocculus during pursuit eye movements. Neurosci. Lett. 72, 163–168 (1986).

27. Krauzlis, R. J. & Lisberger, S. G. Simple spike responses of gaze velocity Purkinje cells in the floccular lobe of the monkey during the onset and offset of pursuit eye movements. J. Neurophysiol. 72, 2045–2050 (1994).

28. Lisberger, S. G., Pavelko, T. A., Bronte-Stewart, H. M. & Stone, L. S. Neural basis for motor learning in the vestibuloocular reflex of primates. II. Changes in the responses of horizontal gaze velocity Purkinje cells in the cerebellar flocculus and ventral paraflocculus. J. Neurophysiol. 72, 954–973 (1994).

29. Raghavan, R. T. & Lisberger, S. G. Responses of Purkinje cells in the oculomotor vermis of monkeys during smooth pursuit eye movements and saccades: comparison with floccular complex. J. Neurophysiol. jn.00209.2017 (2017) doi:10.1152/jn.00209.2017.

30. Medina, J. F. & Lisberger, S. G. Encoding and decoding of learned smooth-pursuit eye movements in the floccular complex of the monkey cerebellum. J. Neurophysiol. 102, 2039–2054 (2009).

31. Lima, S., Hromádka, T., Znamenskiy, P. & Zador, A. PINP: a new method of tagging neuronal populations for identification during in vivo electrophysiological recording. PloS One 4, (2009).

32. Hensbroek, R. A. et al. Identifying Purkinje cells using only their spontaneous simple spike activity. J. Neurosci. Methods 232, 173–180 (2014).

33. Ruigrok, T. J. H., Hensbroek, R. A. & Simpson, J. I. Spontaneous Activity Signatures of Morphologically Identified Interneurons in the Vestibulocerebellum. J. Neurosci. 31, 712–724 (2011).

34. Van Dijck, G. et al. Probabilistic Identification of Cerebellar Cortical Neurones across Species. PLOS ONE 8, e57669 (2013).

35. Lisberger, S. G. & Fuchs, A. F. Role of primate flocculus during rapid behavioral modification of vestibuloocular reflex. I. Purkinje cell activity during visually guided horizontal smooth-pursuit eye movements and passive head rotation. J. Neurophysiol. 41, 733–763 (1978).

36. Shinomoto, S., Shima, K. & Tanji, J. Differences in Spiking Patterns Among Cortical Neurons. Neural Comput. 15, 2823–2842 (2003).

37. Rambold, H., Churchland, A., Selig, Y., Jasmin, L. & Lisberger, S. G. Partial Ablations of the Flocculus and Ventral Paraflocculus in Monkeys Cause Linked Deficits in Smooth Pursuit Eye Movements and Adaptive Modification of the VOR. J. Neurophysiol. 87, 912–924 (2002).

38. Lee, K., Carr, N., Perliss, A. & Chandrasekaran, C. WaveMAP for identifying putative cell types from in vivo electrophysiology. STAR Protoc. 4, 102320 (2023).

39. Lee, E. K. et al. PhysMAP - interpretable in vivo neuronal cell type identification using multi-modal analysis of electrophysiological data. 2024.02.28.582461 Preprint at 10.1101/2024.02.28.582461 (2024).

40. Lee, E. K. et al. Non-linear dimensionality reduction on extracellular waveforms reveals cell type diversity in premotor cortex. eLife 10, e67490 (2021).

41. Bell, C. C. & Grimm, R. J. Discharge properties of Purkinje cells recorded on single and double microelectrodes. J. Neurophysiol. 32, 1044–1055 (1969).

42. Bloedel, J. R. & Roberts, W. J. Action of climbing fibers in cerebellar cortex of the cat. J. Neurophysiol. 34, 17–31 (1971).

43. Mathy, A. et al. Encoding of Oscillations by Axonal Bursts in Inferior Olive Neurons. Neuron 62, 388–399 (2009).

44. Davis, Z. W., Dotson, N. M., Franken, T. P., Muller, L. & Reynolds, J. H. Spike-phase coupling patterns reveal laminar identity in primate cortex. eLife 12, e84512 (2023).

45. Wójcik, D. K. Current Source Density (CSD) Analysis. in Encyclopedia of Computational Neuroscience (eds. Jaeger, D. & Jung, R.) 1–10 (Springer, New York, NY, 2013). doi:10.1007/978-1-4614-7320-6_544-1.

46. Mitzdorf, U. & Singer, W. Excitatory synaptic ensemble properties in the visual cortex of the macaque monkey: A current source density analysis of electrically evoked potentials. J. Comp. Neurol. 187, 71–83 (1979).

47. Mitzdorf, U. Current source-density method and application in cat cerebral cortex: investigation of evoked potentials and EEG phenomena. Physiol. Rev. 65, 37–100 (1985).

48. Mitzdorf, U. Properties of the evoked potential generators: current source-density analysis of visually evoked potentials in the cat cortex. Int. J. Neurosci. 33, 33–59 (1987).

49. Schroeder, C. E., Tenke, C. E., Givre, S. J., Arezzo, J. C. & Vaughan, H. G. Striate cortical contribution to the surface-recorded pattern-reversal vep in the alert monkey. Vision Res. 31, 1143–1157 (1991).

50. Steinschneider, M. et al. Cellular generators of the cortical auditory evoked potential initial component. Electroencephalogr. Clin. Neurophysiol. Potentials Sect. 84, 196–200 (1992).

51. Tahon, K., Wijnants, M., De Schutter, E. & Maex, R. Current source density correlates of cerebellar Golgi and Purkinje cell responses to tactile input. J. Neurophysiol. 105, 1327–1341 (2011).

52. Mendoza-Halliday, D. et al. A ubiquitous spectrolaminar motif of local field potential power across the primate cortex. Nat. Neurosci. 27, 547–560 (2024).

53. Lackey, E. P. et al. Specialized connectivity of molecular layer interneuron subtypes leads to disinhibition and synchronous inhibition of cerebellar Purkinje cells. Neuron 0, (2024).

54. Kole, M. H. P., Letzkus, J. J. & Stuart, G. J. Axon Initial Segment Kv1 Channels Control Axonal Action Potential Waveform and Synaptic Efficacy. Neuron 55, 633–647 (2007).

55. Walsh, J. V., Houk, J. C., Atluri, R. L. & Mugnaini, E. Synaptic Transmission at Single Glomeruli in the Turtle Cerebellum. Science 178, 881–883 (1972).

56. Heine, S. A., Highstein, S. M. & Blazquez, P. M. Golgi Cells Operate as State-Specific Temporal Filters at the Input Stage of the Cerebellar Cortex. J. Neurosci. 30, 17004–17014 (2010).

57. Vos, B. P., Volny-Luraghi, A. & De Schutter, E. Cerebellar Golgi cells in the rat: receptive fields and timing of responses to facial stimulation. Eur. J. Neurosci. 11, 2621–2634 (1999).

58. Simpson, J. I., Hulscher, H. C., Sabel-Goedknegt, E. & Ruigrok, T. J. H. Between in and out: linking morphology and physiology of cerebellar cortical interneurons. in Progress in Brain Research vol. 148 329–340 (Elsevier, 2005).

59. Holtzman, T., Rajapaksa, T., Mostofi, A. & Edgley, S. A. Different responses of rat cerebellar Purkinje cells and Golgi cells evoked by widespread convergent sensory inputs. J. Physiol. 574, 491–507 (2006).

60. Edgley, S. A. & Lidierth, M. The discharges of cerebellar Golgi cells during locomotion in the cat. J. Physiol. 392, 315–332 (1987).

61. Dugué, G. P. et al. Electrical Coupling Mediates Tunable Low-Frequency Oscillations and Resonance in the Cerebellar Golgi Cell Network. Neuron 61, 126–139 (2009).

62. Vervaeke, K. et al. Rapid Desynchronization of an Electrically Coupled Interneuron Network with Sparse Excitatory Synaptic Input. Neuron 67, 435–451 (2010).

63. Hensbroek, R. A., Ruigrok, T. J. H., van Beugen, B. J., Maruta, J. & Simpson, J. I. Visuo-Vestibular Information Processing by Unipolar Brush Cells in the Rabbit Flocculus. The Cerebellum 14, 578–583 (2015).

64. Mugnaini, E., Sekerková, G. & Martina, M. The unipolar brush cell: A remarkable neuron finally receiving deserved attention. Brain Res. Rev. 66, 220–245 (2011).

65. Guo, C., Huson, V., Macosko, E. Z. & Regehr, W. G. Graded heterogeneity of metabotropic signaling underlies a continuum of cell-intrinsic temporal responses in unipolar brush cells. Nat. Commun. 12, 5491 (2021).

66. Haar, S., Givon-Mayo, R., Barmack, N. H., Yakhnitsa, V. & Donchin, O. Spontaneous Activity Does Not Predict Morphological Type in Cerebellar Interneurons. J. Neurosci. 35, 1432–1442 (2015).

67. Sedaghat-Nejad, E. et al. P-sort: an open-source software for cerebellar neurophysiology. J. Neurophysiol. 126, 1055–1075 (2021).

68. Markanday, A. et al. Using deep neural networks to detect complex spikes of cerebellar Purkinje cells. J. Neurophysiol. 123, 2217–2234 (2020).

69. Zur, G. & Joshua, M. Using extracellular low frequency signals to improve the spike sorting of cerebellar complex spikes. J. Neurosci. Methods 328, 108423 (2019).

70. Higgins, I. et al. beta-VAE: Learning Basic Visual Concepts with a Constrained Variational Framework. in (2016).

71. Kingma, D. P. & Welling, M. Auto-Encoding Variational Bayes. Preprint at 10.48550/arXiv.1312.6114 (2022).

72. Lindén, H., Pettersen, K. H. & Einevoll, G. T. Intrinsic dendritic filtering gives low-pass power spectra of local field potentials. J. Comput. Neurosci. 29, 423–444 (2010).

73. Teleńczuk, B. et al. Local field potentials primarily reflect inhibitory neuron activity in human and monkey cortex. Sci. Rep. 7, 40211 (2017).

74. Lainé, J. & Axelrad, H. Extending the cerebellar Lugaro cell class. Neuroscience 115, 363–374 (2002).

75. Lainé, J. & Axelrad, H. The candelabrum cell: A new interneuron in the cerebellar cortex. J. Comp. Neurol. 339, 159–173 (1994).

76. Osorno, T. et al. Candelabrum cells are ubiquitous cerebellar cortex interneurons with specialized circuit properties. Nat. Neurosci. 25, 702–713 (2022).

77. Mountcastle, V. B., Talbot, W. H., Sakata, H. & Hyvärinen, J. Cortical neuronal mechanisms in flutter-vibration studied in unanesthetized monkeys. Neuronal periodicity and frequency discrimination. J. Neurophysiol. 32, 452–484 (1969).

78. Johnston, K., DeSouza, J. F. X. & Everling, S. Monkey Prefrontal Cortical Pyramidal and Putative Interneurons Exhibit Differential Patterns of Activity Between Prosaccade and Antisaccade Tasks. J. Neurosci. 29, 5516–5524 (2009).

79. Ardid, S. et al. Mapping of Functionally Characterized Cell Classes onto Canonical Circuit Operations in Primate Prefrontal Cortex. J. Neurosci. 35, 2975–2991 (2015).

80. Trainito, C., von Nicolai, C., Miller, E. K. & Siegel, M. Extracellular Spike Waveform Dissociates Four Functionally Distinct Cell Classes in Primate Cortex. Curr. Biol. 29, 2973–2982.e5 (2019).

81. Ramachandran, R. & Lisberger, S. G. Normal Performance and Expression of Learning in the Vestibulo-Ocular Reflex (VOR) at High Frequencies. J. Neurophysiol. 93, 2028–2038 (2005).

82. Robinson, D. A. A Method of Measuring Eye Movement Using a Scleral Search Coil in a Magnetic Field. IEEE Trans. Bio-Med. Electron. 10, 137–145 (1963).

83. Rashbass, C. The relationship between saccadic and smooth tracking eye movements. J. Physiol. 159, 326–338 (1961).

84. Carl, J. R. & Gellman, R. S. Human smooth pursuit: stimulus-dependent responses. J. Neurophysiol. 57, 1446–1463 (1987).

85. Hall, N. J., Herzfeld, D. J. & Lisberger, S. G. Evaluation and resolution of many challenges of neural spike sorting: a new sorter. J. Neurophysiol. 126, 2065–2090 (2021).

86. Lisberger, S. G. & Pavelko, T. A. Vestibular signals carried by pathways subserving plasticity of the vestibulo-ocular reflex in monkeys. J. Neurosci. 6, 346–354 (1986).

87. Beau, M., et al. NeuroPyxels: loading, processing and plotting Neuropixels data in python. Zenodo 10.5281/zenodo.5509776 (2021).

88. Mitra, P. P. & Pesaran, B. Analysis of dynamic brain imaging data. Biophys. J. 76, 691–708 (1999).

89. Kingma, D. P., Rezende, D. J., Mohamed, S. & Welling, M. Semi-supervised learning with deep generative models. in Proceedings of the 27th International Conference on Neural Information Processing Systems - Volume 2 3581–3589 (MIT Press, Cambridge, MA, USA, 2014). 10.48550/arXiv.1406.5298.

90. Kingma, D. P. & Ba, J. Adam: A Method for Stochastic Optimization. Preprint at 10.48550/arXiv.1412.6980 (2017).

91. Loshchilov, I. & Hutter, F. SGDR: Stochastic Gradient Descent with Warm Restarts. in (2016).

92. Fukushima, K. Visual Feature Extraction by a Multilayered Network of Analog Threshold Elements. IEEE Trans. Syst. Sci. Cybern. 5, 322–333 (1969).

93. Ganaie, M. A., Hu, M., Malik, A. K., Tanveer, M. & Suganthan, P. N. Ensemble deep learning: A review. Eng. Appl. Artif. Intell. 115, 105151 (2022).

